# Phytochemical Characterization of Bio-active Compounds in Hydroethanolic Extract of *Elaeocarpus ganitrus* leaves using HPLC, LC-MS, and HPTLC Analyses

**DOI:** 10.1101/2023.10.12.562125

**Authors:** Jyotsana Khushwaha, Alpana Joshi, Shiva Sharma, Subrata K. Das

## Abstract

Bioactive compounds have various applications in different industries, including food, pharmaceutical, and cosmetic industries, demonstrating the need to identify the best-standardized technique to screen the phytochemical profile of medicinal plants. This study aimed to characterize the bioactive compounds in the hydroethanolic extracts of *Elaeocarpus ganitrus* leaves using various analytical techniques: HPLC, LC-MS, and HPTLC. Air-dried leaves of *E. ganitrus* were extracted with 70% ethanol. The phytochemical composition of crude extracts was analyzed by the High performance liquid chromatography (HPLC) method, and a total of 93 compounds, including 46 flavonoids, 17 phenols, 14 polyphenols, 3 phenolic acid, 3 phenolic glycosides, 2 flavonoid glycosides, 2 glycosides, 2 phenylpropanoid glycoside, 1 hydroxycinnamic acid, 1 lignan, 1 tannin, and 1 terpene glycoside were detected and quantified. The Liquid chromatography mass spectrometry (LC-MS) analyses identified 11 major eleven compounds: quercetin (803.0215 µg/L), gallic acid (726.13 µg/L), ferulic acid (652.34 µg/L), chlorogenic acid (651.021µg/L), pinocembrin (264.11 µg/L), p-aminobenzoic acid (251.021 µg/L), epicatechin (246.02 µg/L), catechin (161.51 µg/L), caffeic acid (123.31 µg/L), syringaldehyde (116.31 µg/L), and naringenin (106.31 µg/L). The chemical fingerprinting was carried out by high performance thin layer chromatography (HPTLC), and HPTLC fingerprint qualitatively revealed a predominant amount of gallic acid (48.64 %), curcumin (15.21 %), caffeic acid (12.19 %) and cinnamic acid (6.50 %). A significant amount of bioactive constituents in a hydroethanolic extract of *E. ganitrus* leaves indicates the plant’s therapeutic potential, including antioxidant, anti-inflammatory, antidiabetic, anticancer, neuroprotective, and cardio-protective activities.

## 1. Introduction

Medicinal plants constitute the basis of traditional and modern primary healthcare. Over 80% of the population, mainly of developing countries, depend on traditional and herbal medicine. In the past two decades, there has been substantial growth in the use of medicinal plants to prevent disease and promote health. The current pharmacopeia contains at least 25% plant-based medications; however, the medicinal plants must meet quality, safety, and efficacy standards for proper utilization. One of the most significant difficulties related to quality is that commercial medicinal plants are available in powdered form, making it challenging to identify specific plant parts or plant species **(Salmerón-Manzano *et al*., 2020)**.

*Elaeocarpus ganitrus* (Rudraksha) belongs to the family Elaeocarpaceae and has been well-known from ancient times for its medicinal importance **(Kumari *et al*., 2018; Rai *et al*., 2018; Sharma *et al*., 2022; Joshi and Kushwaha, 2023)**. These fruits are commonly found in India, specifically in the Himalayan and Gangetic plain regions, Nepal, Indonesia, and Java. The pharmacological action of *Elaeocarpus sp.* is due to the presence of bioactive phytochemicals and numerous studies revealed that petroleum ether, ethanol, and water extracts of *Elaeocarpus sp.* contain several alkaloids (elaeocarpidine, elaeocarpine, rudrakine), polyphenols (flavonoids, quercetin, tannin), phytosterols, fat, carbohydrates, proteins, gallic and ellagic acid **(Johns and Lamberton, 1973; Katavic *et al*., 2007; Sudrajat and Timotius, 2022)**. The major identified biochemical compounds are isoelaeocarpine, epiisoelaeocarpiline, epielaeocarpiline, alloelaeocarpiline, and pseudo-epiiso-elaeocarpilline **(Johns *et al*., 1970; Katavic *et al*., 2006; Katavic *et al*., 2007; Ezeoke *et al*., 2018; Sudrajat and Timotius, 2022)**.

*E. ganitrus* beads (Rudraksha) are known for their therapeutic potential against several disorders like stress, anxiety, insomnia, skin diseases, leprosy, hysteria, hyperglycemia, coma, leucorrhoea, infertility, asthma, hypertension, diabetes, arthritis, rheumatism, cardiovascular and liver diseases (**Rai *et al*., 2018; Sharma *et al*., 2022)**. There are evidences in literature indicating their sedative, analgesic, anticonvulsant, anti-inflammatory, antioxidant, antiepileptic, hypnotic, antipyretic, antihypertensive, antidiabetic, antimicrobial, anxiolytic, anti-cancerous, anti-asthmatic, nephroprotective, immune-stimulator, and electromagnetic properties **(Ray *et al*., 1979; Fang *et al*., 1984; Ito *et al*., 2002; Katavic *et al*., 2006; Katavic *et al*., 2007; Meng *et al*., 2008; Shitamoto *et al*., 2010; Pan *et al*., 2012; Bordoloi *et al*., 2017; Liyanaarachchi *et al*., 2018; Kim *et al*., 2018; Ezeoke *et al*., 2018; Hong *et al*., 2019; Ogundele and Das, 2019; Turner *et al*., 2020; Ogundele *et al*., 2021; Kim *et al*., 2021; Banerjee *et al*., 2022; Joo *et al*., 2022**). A summary of phytochemical investigations on *Elaeocarpus* species is listed in **Table 1**.

**Table 1.** Literature survey on phytochemical profile for *Elaeocarpus* species.

| Name of the species | Phytochemicals | References |
| --- | --- | --- |
| *E. angustifolius* | (±)-8,9-Dehydroelaecarpine, (±)-Elaecarpine trifluoroacetate, (±)- 9-Epielaecarpine cis-N-oxide trifluoroacetate | Hong *et al.*, 2019 |
| *E. chinensis* | Cucurbitacins D, Elaecarpucins A-H | Pan *et al.*, 2012 |
| *E. dolichostylus* | Cucurbitacin F | Fang *et al.*, 1984 |
| *E. flooribundus* | Gallic acid, myricitrin, mearnsitrin, myricetin, and mearnsetin, Phytol, a-tocopherolquinone, Euphorbol, Phaeophytins | Ogundele and Das, 2019; Ogundele *et al.*, 2021, Banerjee *et al.*, 2022 |
| *E. fucoides* | Elaecarpine, Elaecarpenine, Isoelaecarpine, | Katavic *et al.*, 2007 |
| *E. ganitrus* | Rudrakine | Ray *et al.*, 1979 |
| *E. grandis* | (-)-Isoelaecarpiline, Grandisine C, D, E, G, | Katavic *et al.*, 2006 |
| *E. hainanensis* | Cucurbitacins D | Meng *et al.*, 2008 |
| *E. japonicus* | Elaecarpionoside | Shitamoto *et al.*, 2010 |
| *E. lanceofolius* | Myricetin, 4'-Methylmyricetin, Mearnsetin, Triacontanoic acid, Octatriacontan-1-ol, Dotriacontane | Ray *et al.*, 1979; Bordoloi *et al.*, 2017 |
| *E. mastersii* | Cucurbitacins D, Cucurbitacin F, 4'-O-Methylellagic acid 3-(2'',3''-di-O-acetyl)-α-l-rhamnoside, 4,4'-O-Dimethylellagic acid 3-(2'',3''-di-O-acetyl)-α-l-rhamnoside | Ito *et al.*, 2002 |
| *E. reticulatus* | Cucurbitacin-I, Proanthocyanidins anthocyanins, | Turner *et al.*, 2020 |
| *E. serratus* | Dibutyl succinate, Phytosterol, Elastase, Hyaluronidase, Tyrosinase | Liyanaarachchi *et al.*, 2018 |
| *E. sylvestris* | Geraniin, 1, 2, 3, 4, 6-penta-O-galloyl-β-D-glucose (PGG), elaeocarpusin, | Kim *et al.*, 2018;<br>Kim *et al.*, 2021;<br>Joo *et al.*, 2022 |
| *E. tectorius* | Tectoricine, Tectoraline, Tectoramidines A, B | Ezeoke *et al.*, 2018 |

The plant contains abundant bioactive compounds in different concentrations and polarity. A key challenge in screening plant phytochemical profiles is extraction and characterization methods. The combination of different analytical techniques, such as High-performance liquid chromatography (HPLC), Liquid chromatography-mass spectrometry (LC-MS), and high-performance thin-layer chromatography (HPTLC), can be applied to detect bioactive constituents in plant extracts. These analytical techniques are effective for ensuring the quality of raw plant material and can be used to analyze various plant extracts **(Nile and Park, 2014)**. The phytochemical profile of the ethanolic fraction of *E. floribundus* fruits displayed various biological activities, including antimicrobial **(Sircar *et al*., 2017)**. HPLC and GC-MS analyses was conducted out to examine the bioactive constituents present in the fruits of *E. oblongus, E. serratus,* and *E. tectorius* **(Muthuswamya and Senthamarai, 2014; Mundaragi *et al*., 2019; de Lima *et al*., 2019)**. LC-MS combines the separation abilities of liquid chromatography against a target compound. LC-MS profile of *E. grandiflorus* and *E. sphaericus* demonstrated the bioactive compounds significantly **(Primiani *et al*., 2021; Habibah *et al*., 2021)**.

However, the phytochemical profiling of *E. ganitrus* using HPLC, LC-MS, and HPTLC has not been reported. Hence, the present investigation aimed to conduct a qualitative and quantitative assessment of phytochemical constituents in the hydroethanolic extract of *E. ganitrus* leaves using three different analytical techniques: High-performance liquid chromatography (HPLC), Liquid chromatography-mass spectrometry (LC-MS), and high-performance thin layer chromatography (HPTLC).

## 2. Material and Methods

### 2.1. Sample collection and hydroethanolic extract preparation

Fresh leaves samples of *E. ganitrus* were harvested at the Shobhit Institute of Engineering & Technology (Deemed-to-be-University), Modipuram, Meerut, India, with the coordinates of the sites, Latitude 29.071274° and Longitude 77.711929°. Leaf samples were rinsed with double distilled water and air dried under shade conditions until all moisture content was gone. The plant samples (2 g) were ground into a fine powder using liquid nitrogen by mortar and pestle. The hydroethanolic extract was prepared by adding 70 % ethanol (10 ml) and incubated for 1 week at room temperature (**Figure 1**). Following centrifugation and filtration, extracts were lyophilized and stored at -80 ^◦^C.

**Figure 1.**
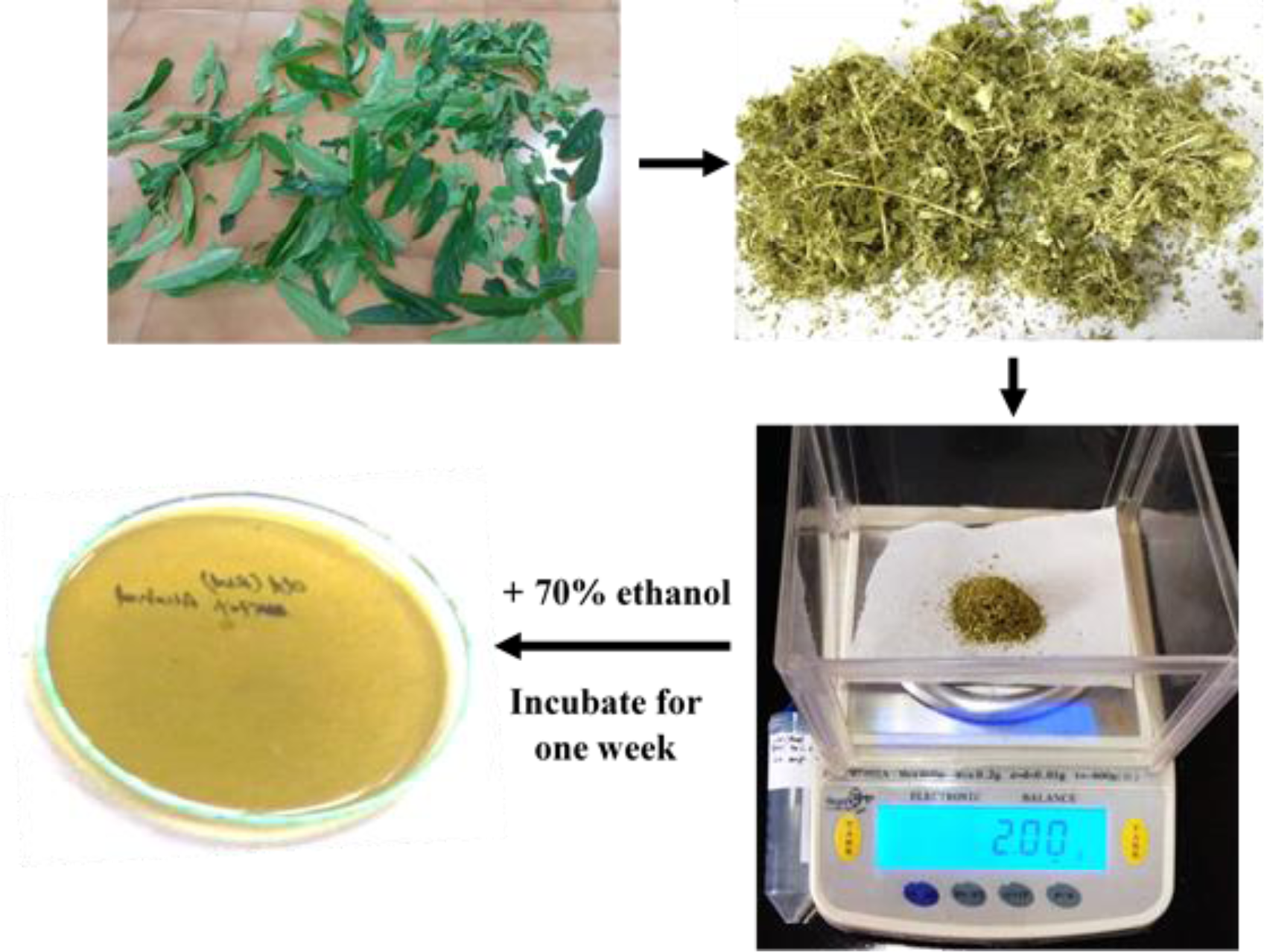
The process of hydroethanolic extract preparation from *E. ganitrus* leaves.

### 2.2. High-Performance Liquid Chromatography (HPLC) analysis

Reagents of analytical grade Toluene, Ethyl acetate, Formic acid, Gallic acid, Catechin, Caffeic acid, Berberine, Rutin, Cinnamic acid, and Curcumin were obtained (Sigma-Aldrich, India). Precoated TLC Aluminum sheets silica gel 60F254 (10 x 10 cm, 0.2 mm thick) were obtained from E. Merck Ltd, Mumbai. The extract was diluted in 50 % methanol (1 mg/ml) and subjected to HPLC analysis. Waters binary HPLC system (Waters Corporation, Milford, MA, USA), equipped with column oven, auto-sampler (Waters 2707), and photodiode array (PDA) detector (Waters 2998), was used for the analyses. A reversed-phase C18 analytical column (4.60×250 mm, 5 μm particle size; Sunfire, Waters, U.S.A.) was utilized at 30 °C column temperature. Binary gradient was used with 0.1% HCOOH in water (A) and Acetonitrile (B), and a run time of 35 minutes at a flow rate of 1ml/min was used for the analysis. The injection volume was 20 µl (E. ganitrus leaf extracts) and 10 µl (standard mix) at different concentrations. The identities of constituents were also confirmed with a photodiode array (PDA) detector by comparison with ultraviolet (UV) spectra of standards in the wavelength at 280 nm and 325 nm.

### 2.3. LC-MS Analysis

The hydroethanolic extract of *E. ganitrus* leaves was used for LC-MS analysis **(Singh *et al*., 2022)**. The LC/MS instrument is equipped with an Electron Spray Ionization (ESI) ion source operating in a positive and negative ion mode. The capillary temperature was kept at 280 °C, and the sample flow rate was 8 μL/min. A mass range was selected from 50-1000 Da with a scanning time of 0.2 s. The elution was carried out using 156 gradient elution of 0.1 % formic acid in water (solvent A) and 0.1 % formic acid in 157 acetonitrile (solvent B) with a 400 µl/min flow rate. The solvent gradient program was started with 95-90 % of the mobile phase for 0-5 min, 90-80 % for 5-10 min, 80-60 % for 10-20 min, 60-40 % for 20-30 min, 40 % for 30-45 min, 40-95 % for 45-46 min, followed by 95 % for 46-50 min. Two microlitres of the test solution were used for screening, and the chromatograph was continuously tracked for 45 minutes.

### 2.4. HPTLC Analysis

Analysis was performed on a Camag HPTLC system equipped with a sample applicator ATS4, ADC2 development chamber, and TLC Scanner; TLC Visualizer and WinCats integration software were used. The standard solutions of gallic acid, catechin, caffeic acid, berberine, rutin, colchicine, cinnamic acid, and Curcumin were accurately weighed (10 mg), and the solution was made up to 10 ml with methanol (1 mg/ml). From the stock solution of the standards, 0.1 ml was pipetted out and further diluted up to 1 ml to obtain the final concentration of 100 µg/ml. For standard mixture preparation, different standards were mixed to get a 100 ppm final concentration for all standards in methanol. Hydroethanolic extract of *E. ganitrus* leaves, and the standards were spotted on a precoated TLC Aluminum sheets silica gel 60 F254 (20×10cm, 0.2mm thickness) as 8mm wide bandwidth by using automatic TLC applicator ATS 4,10mm from the bottom. The Mobile phase used was Toluene: Ethyl acetate: Formic acid (5:4:1v/v). The plates were saturated in ADC2 for 20 min. After development, the plates were dried in ADC2 and scanned at 254 nm, 366, and after derivatization at 540 nm using CAMAG Scanner. The plates were photographed at an optimized wavelength of 254 nm, 366 nm, and 540 nm.

## 2.5. Data analysis

The chemical structure for each compound identified in the hydroethanolic extract of *E. ganitrus* leaves using HPLC, LC-MS, and HPTLC was searched using online database software (www.chemspider.com).

## 3. Results & Discussion

### 3.1. Identification and quantification of marker compounds by HPLC

Phytochemical profiling of hydroethanolic extracts of *E. ganitrus* leaves was performed using the HPLC analysis. A binary gradient method for HPLC was developed and optimized. The hydroethanolic extract was analyzed along with the mixture of standard marker compounds. Altogether, 93 phenolic compounds in the leaves of *E. ganitrus* were identified and quantified, including 2 flavonoid glycosides, 46 flavonoids, 2 glycosides, 1 hydroxycinnamic acid, 1 lignan, 3 phenolic acid, 3 phenolic glycosides, 17 phenols, 2 phenylpropanoid glycoside, 14 polyphenols, 1 tannin, and 1 terpene glycoside using HPLC analysis **(Table 2)**. Each compound was identified and confirmed using its retention time (RT) and UV profile in a photodiode array (PDA) detector under similar conditions (**Figure 2** **and** **Figure 3**).

**Figure 2.**
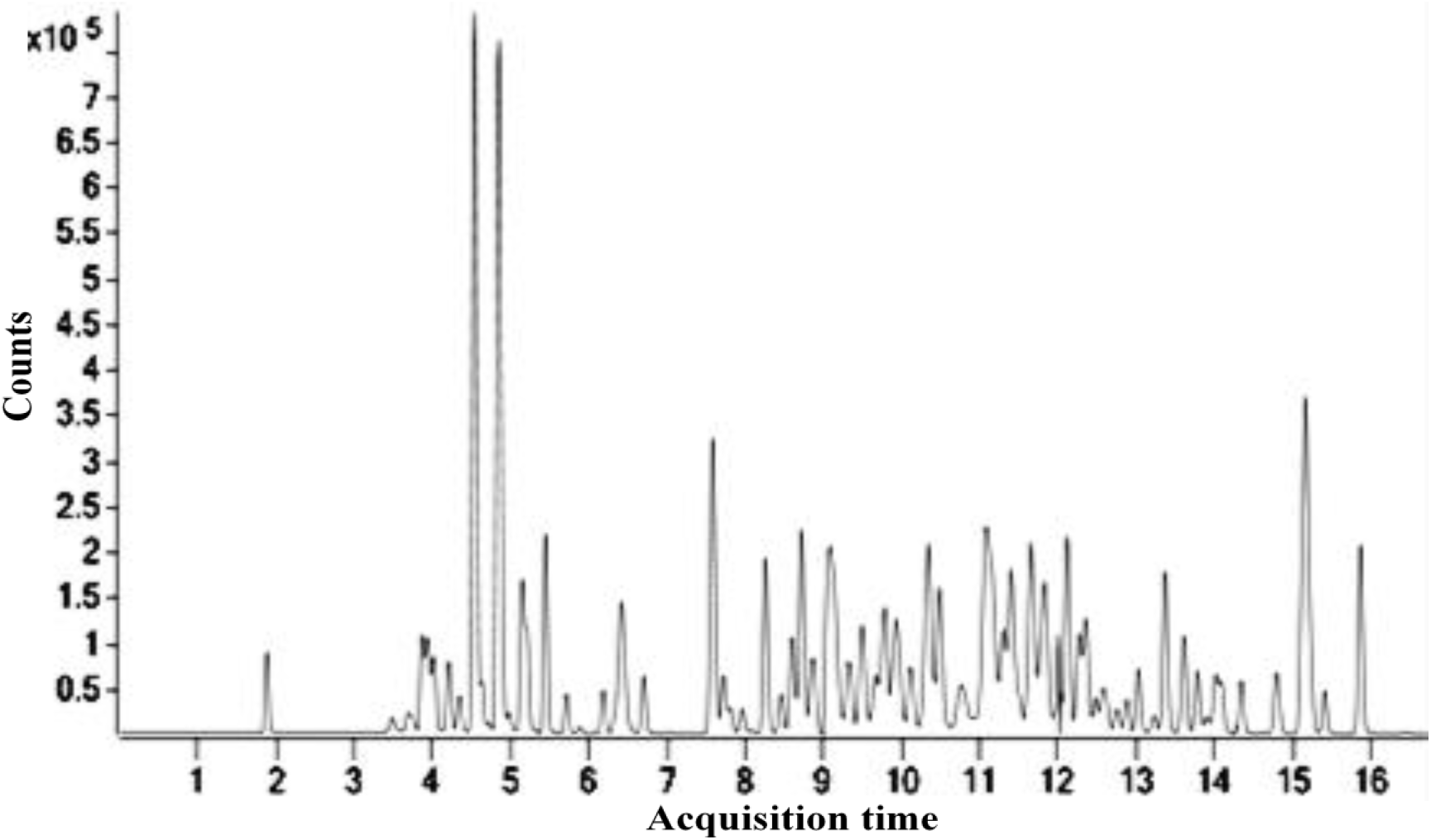
HPLC chromatogram of identified phytochemical constituents’ profile hydroethanolic extract of *E. ganitrus* leaves.

**Figure 3.**
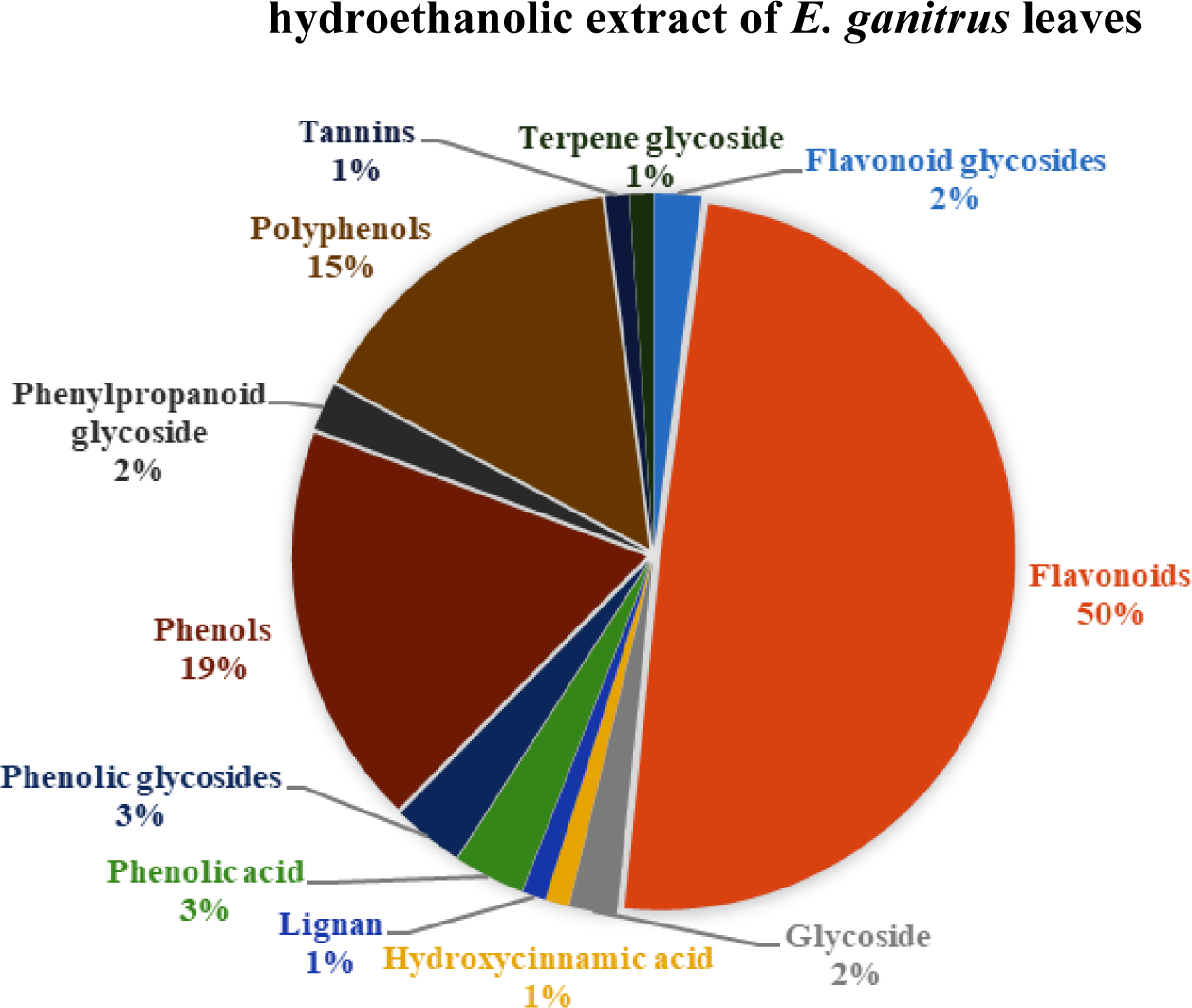
The phytochemical composition identified in HPLC analysis in a hydroethanolic fraction of *E. ganitrus* leaves.

**Table 2.**
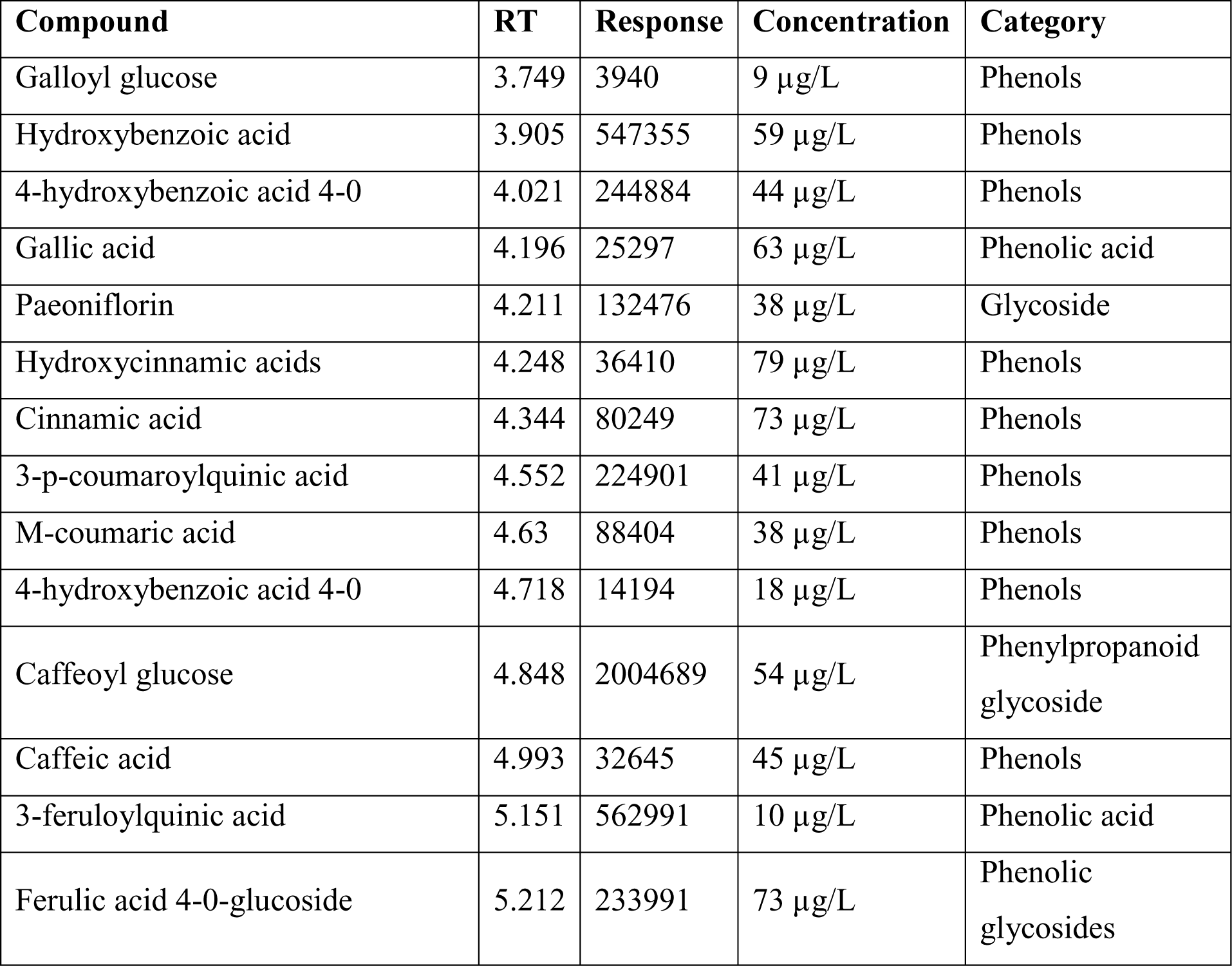
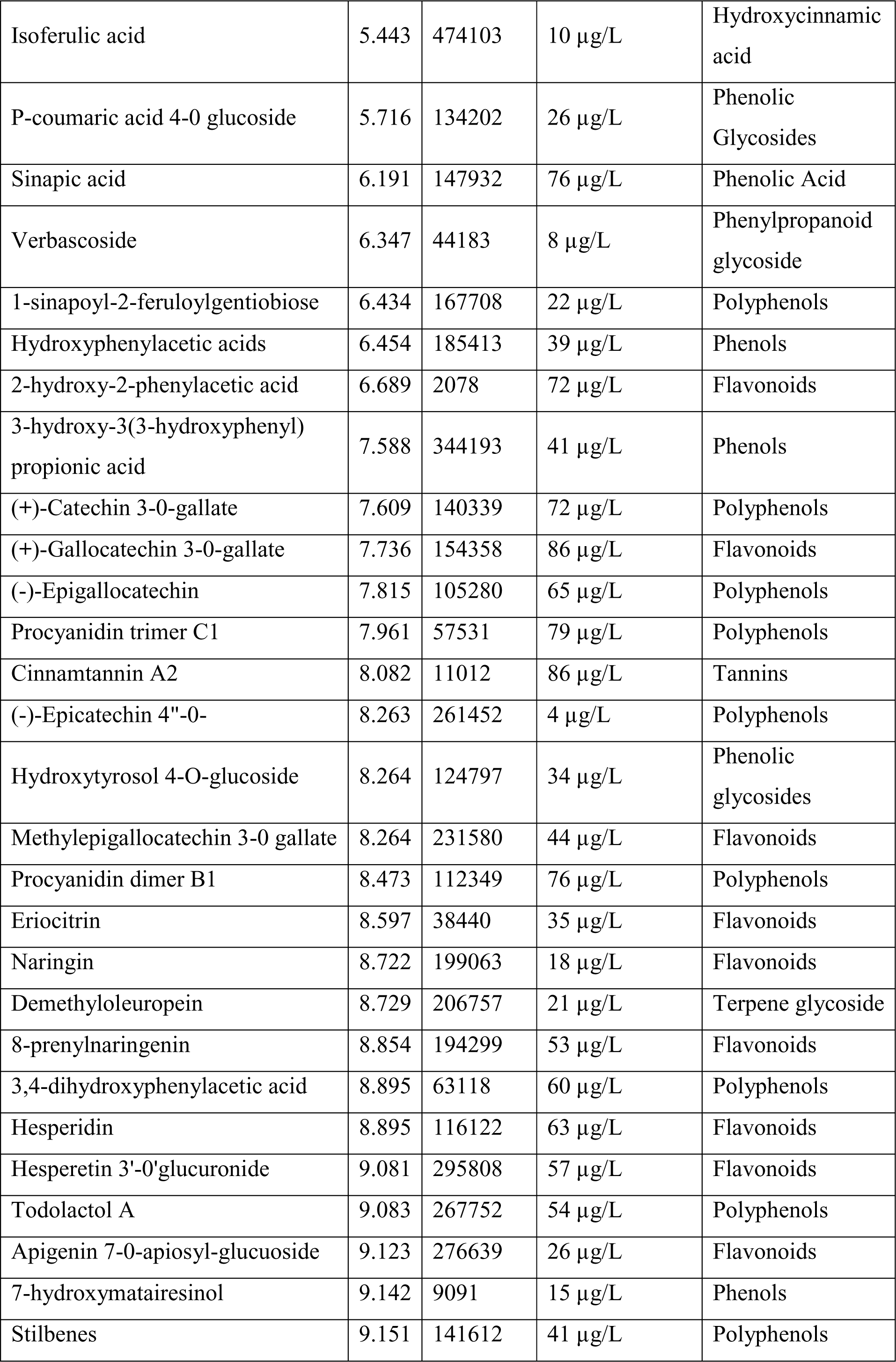
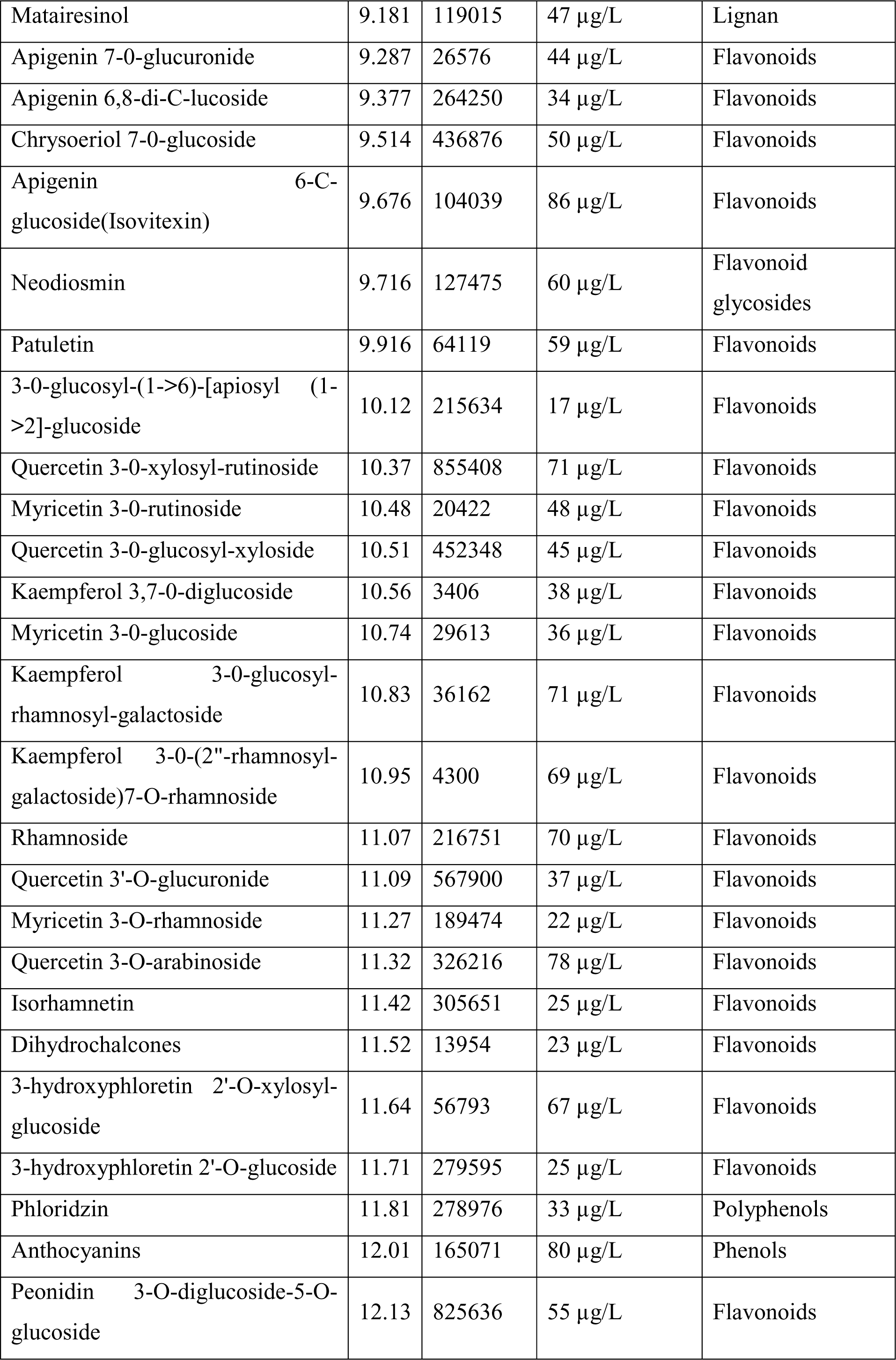
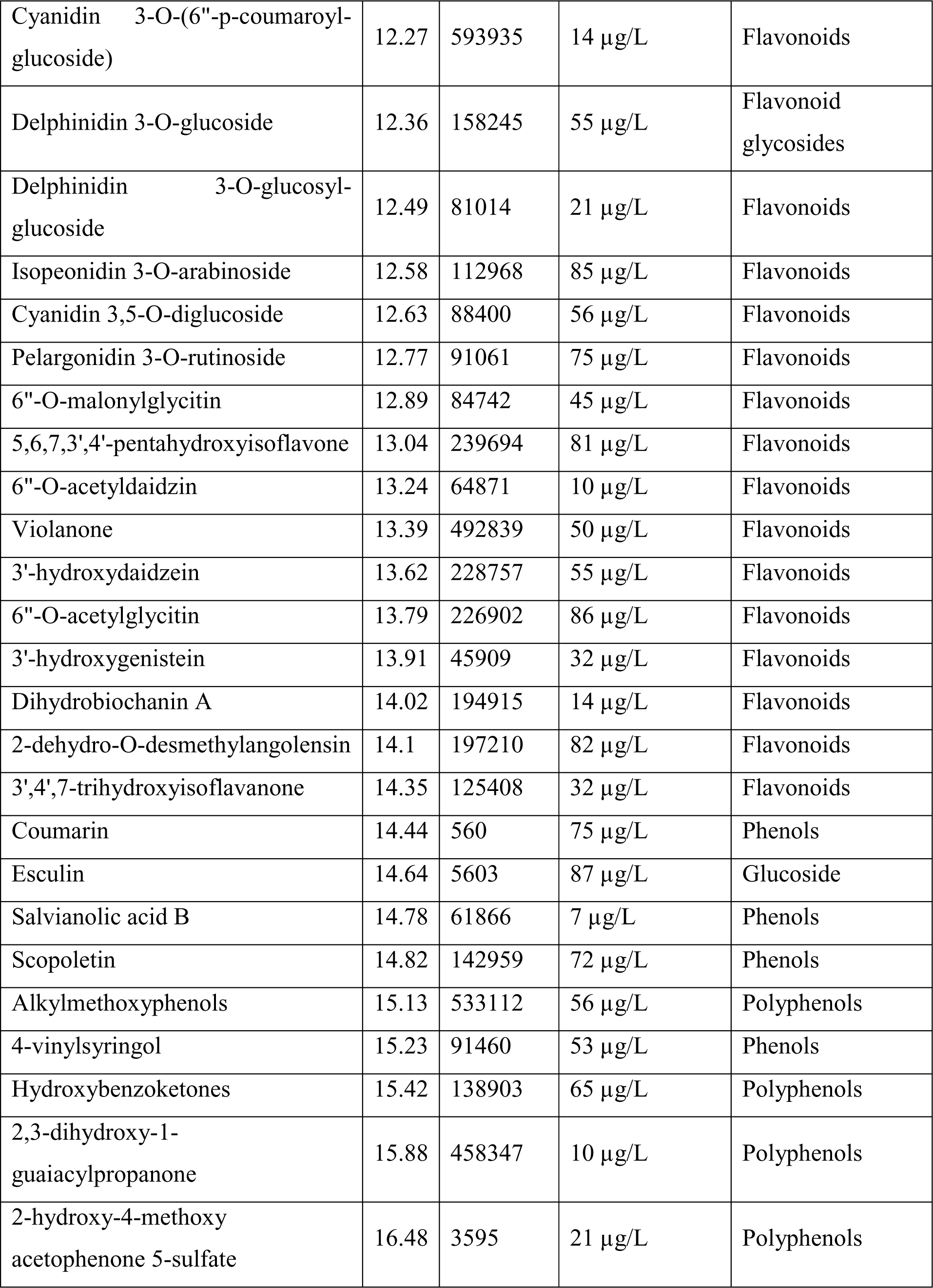
The phytochemical profile of the *E. ganitrus* hydroethanolic extract identified using HPLC.

The flavonoids, including flavanones, flavanols, flavonols, flavones, and isoflavones, were the most abundant compounds annotated in the hydroethanolic fraction of *E. ganitrus* leaves **(Table 2)**. Forty six flavonoids were reported including; 2-hydroxy-2-phenylacetic acid (72 µg/L) at RT 6.689, (+)-Gallocatechin 3-0-gallate (86 µg/L) at RT 7.736, Methylepigallocatechin 3-0 gallate (44 µg/L) at RT 8.264, Eriocitrin (35 µg/L) at RT 8.597, Naringin (18 µg/L) at RT 8.722, 8-prenylnaringenin (53 µg/L) at RT 8.854, Hesperidin (63 µg/L) at RT 8.895, Hesperetin 3’-0’glucuronide (57 µg/L) at RT 9.081, Apigenin 7-0-apiosyl-glucuoside (26 µg/L) at RT 9.123, Apigenin 7-0-glucuronide (44 µg/L) at RT 9.287, Apigenin 6,8-di-C-lucoside (34 µg/L) at RT 9.377, Chrysoeriol 7-0-glucoside (50 µg/L) at RT 9.514, Apigenin 6-C-glucoside(Isovitexin) (86 µg/L) at RT 9.676, Patuletin (59 µg/L) at RT 9.916, 3-0-glucosyl-(1->6)-[apiosyl (1->2]-glucoside (17 µg/L) at RT 10.12, Quercetin 3-0-xylosyl-rutinoside (71 µg/L) at RT 10.37, Myricetin 3-0-rutinoside (48 µg/L) at RT 10.48, Quercetin 3-0-glucosyl-xyloside (45 µg/L) at RT 10.51, Kaempferol 3,7-0-diglucoside (38 µg/L) at RT 10.56, Myricetin 3-0-glucoside (36 µg/L) at RT 10.74, Kaempferol 3-0-glucosyl-rhamnosyl-galactoside (71 µg/L) at RT 10.83, Kaempferol 3-0-(2“-rhamnosyl-galactoside)7-O-rhamnoside (69 µg/L) at RT 10.95, Rhamnoside (70 µg/L) at RT 11.07, Quercetin 3’-O-glucuronide (37 µg/L) at RT 11.09, Myricetin 3-O-rhamnoside (22 µg/L) at RT 11.27, Quercetin 3-O-arabinoside (78 µg/L) at RT 11.32, Isorhamnetin (25 µg/L) at RT 11.42, Dihydrochalcones (23 µg/L) at RT 11.52, 3-hydroxyphloretin 2’-O-xylosyl-glucoside (67 µg/L) at RT 11.64, 3-hydroxyphloretin 2’-O-glucoside (25 µg/L) at RT 11.71, Peonidin 3-O-diglucoside-5-O-glucoside (55 µg/L) at RT 12.13, Cyanidin 3-O-(6“-p-coumaroyl-glucoside) (14 µg/L) at RT 12.27, Delphinidin 3-O-glucosyl-glucoside (21 µg/L) at RT 12.49, Isopeonidin 3-O-arabinoside (85 µg/L) at RT 12.58, Cyanidin 3,5-O-diglucoside (56 µg/L) at RT 12.63, Pelargonidin 3-O-rutinoside (75 µg/L) at RT 12.77, 6“-O-malonylglycitin (45 µg/L) at RT 12.89, 5,6,7,3’,4’-pentahydroxyisoflavone (81 µg/L) at RT 13.04, 6“-O-acetyldaidzin (10 µg/L) at RT 13.24, Violanone (50 µg/L) at RT 13.39,3’-hydroxydaidzein (55 µg/L) at RT 13.62, 6“-O-acetylglycitin (86 µg/L) at RT 13.79, 3’-hydroxygenistein (32 µg/L) at RT 13.91, Dihydrobiochanin A (14 µg/L) at RT 14.02, 2-dehydro-O-desmethylangolensin (82 µg/L) at RT 14.1, and 3’,4’,7-trihydroxyisoflavanone (32 µg/L) at RT 14.35. Various investigations reported the antiviral, anticancer, neuroprotective, and anti-inflammatory activities of flavonoids **(Muhammad *et al*., 2019; Yuan *et al*., 2021; Ortiz *et al*., 2022; Aboulaghras *et al*., 2022; Salehi *et al*., 2020; Ayvaz *et al*., 2022; Patel *et al*., 2023)**.

The second abundant category was phenols, and 17 compounds were recognized including; Galloyl glucose (9 µg/L) at RT 3.749, Hydroxybenzoic acid (59 µg/L) at RT 3.905, 4-hydroxybenzoic acid 4-0 (44 µg/L) at RT 4.021, Hydroxycinnamic acids (79 µg/L) at RT 4.248, Cinnamic acid (73 µg/L) at RT 4.344, 3-p-coumaroylquinic acid (41 µg/L) at RT 4.552, M-coumaric acid (38 µg/L) at RT 4.63, 4-hydroxybenzoic acid 4-0 (18 µg/L) at RT 4.718, Caffeic acid (45 µg/L) at RT 4.993, Hydroxyphenylacetic acids (39 µg/L) at RT 6.454, 3-hydroxy-3(3-hydroxyphenyl) propionic acid (41 µg/L) at RT 7.588, 7-hydroxymatairesinol (15 µg/L) at RT 9.142, Anthocyanins (80 µg/L) at RT 12.01, Coumarin (75 µg/L) at RT 14.44, Salvianolic acid B (7 µg/L) at RT 14.78, Scopoletin (72 µg/L) at RT 14.82, and 4-vinylsyringol (53 µg/L) at RT 15.23. Phenols are known to exhibit various pharmacological activities such as., antioxidant, anti-inflammatory, antimicrobial, anti-adipogenic, antidiabetic anticancer, and neuroprotective **(Cardile *et al*., 2015; Li *et al*., 2020; Kowalska *et al*., 2021)**.

The third abundant category of phytochemicals was polyphenols. The present study detected 14 polyphenols, including 1-sinapoyl-2-feruloylgentiobiose (22 µg/L) at RT 6.434, (+)-Catechin 3-0-gallate (72 µg/L) at RT 7.609, (-)-Epigallocatechin (65 µg/L) at RT 7.815, Procyanidin trimer C1 (79 µg/L) at RT 7.961, (-)-Epicatechin 4“-0-(4 µg/L) at RT 8.263, Procyanidin dimer B1 (76 µg/L) at RT 8.473, 3,4-dihydroxyphenylacetic acid (60 µg/L) at RT 8.895, Todolactol A (54 µg/L) at RT 9.083, Stilbenes (41 µg/L) at RT 9.151, Phloridzin (33 µg/L) at RT 11.81, Alkylmethoxyphenols (56 µg/L) at RT 15.13, Hydroxybenzoketones (65 µg/L) at RT 15.42, 2,3-dihydroxy-1-guaiacylpropanone (10 µg/L) at RT 15.88, and 2-hydroxy-4-methoxyacetophenone 5-sulfate (21 µg/L) at RT 16.48. Polyphenols are secondary metabolites that exhibit multiple pharmacological activities: anti-infectious, anti-inflammatory, cardio-protective, antimicrobial, antiviral, antimutagenic, antihyperglycemic, and anti-allergic **(Rue *et al*., 2018)**. The major bioactive compounds identified by HPLC analysis were displayed with their classification and pharmacological activities **(Table 4)**.

**Table 3:**
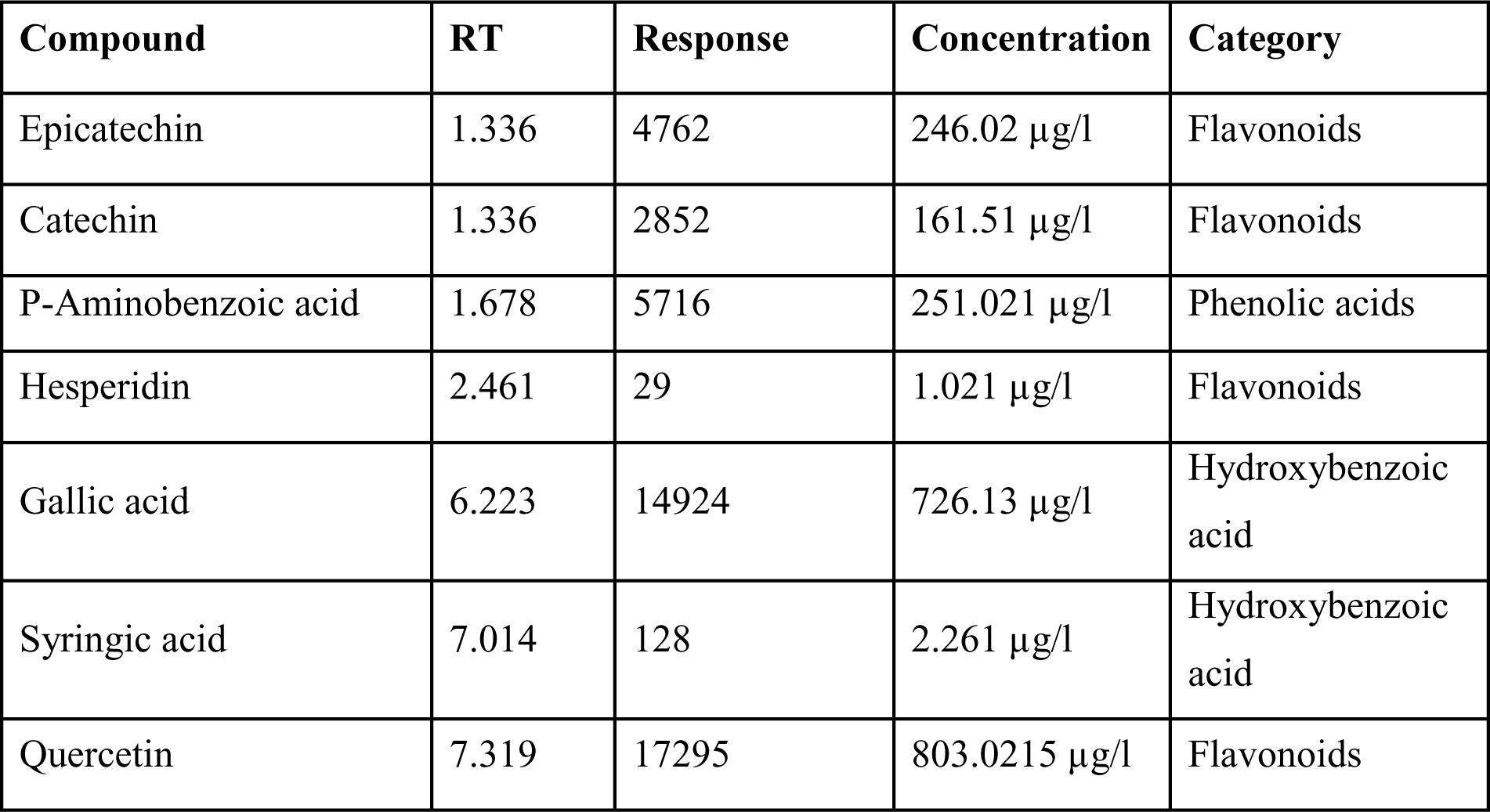
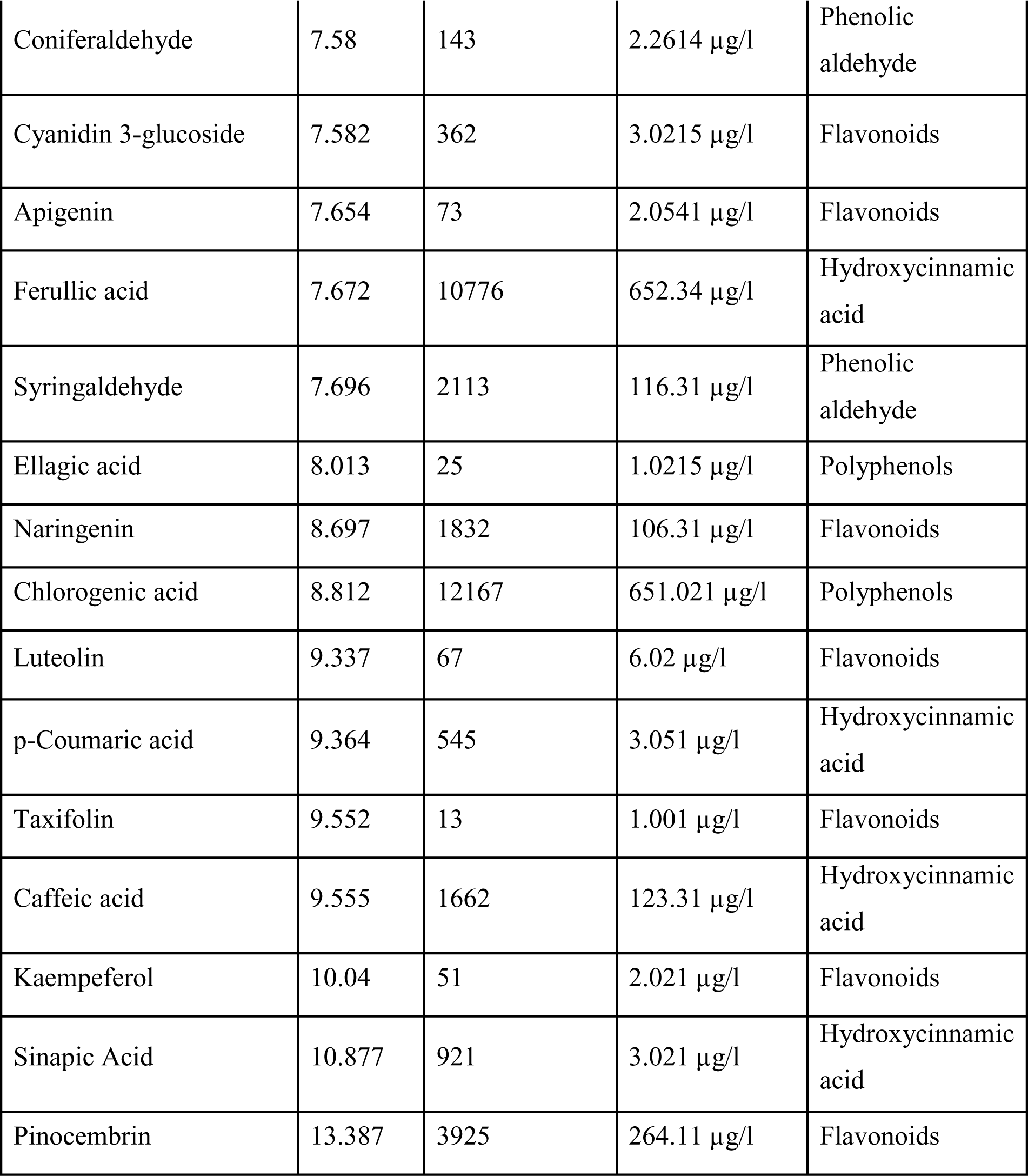
Quantification of phytochemicals in hydroethanolic extract of *E. ganitrus* leaves through LC-MS.

**Table 4:**
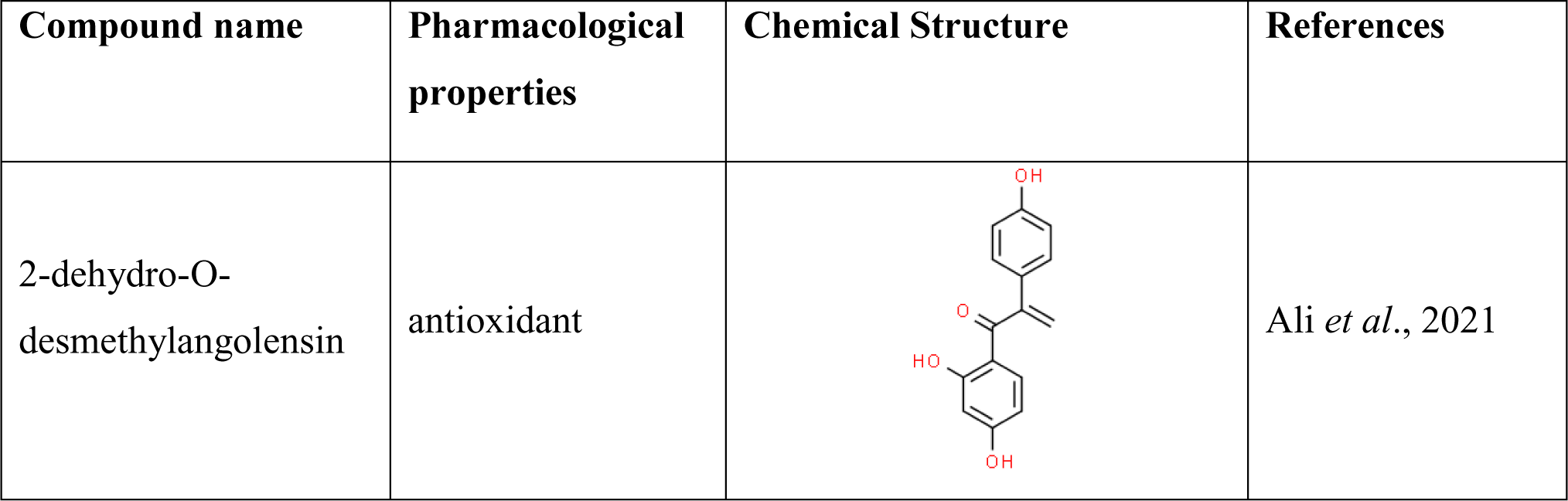
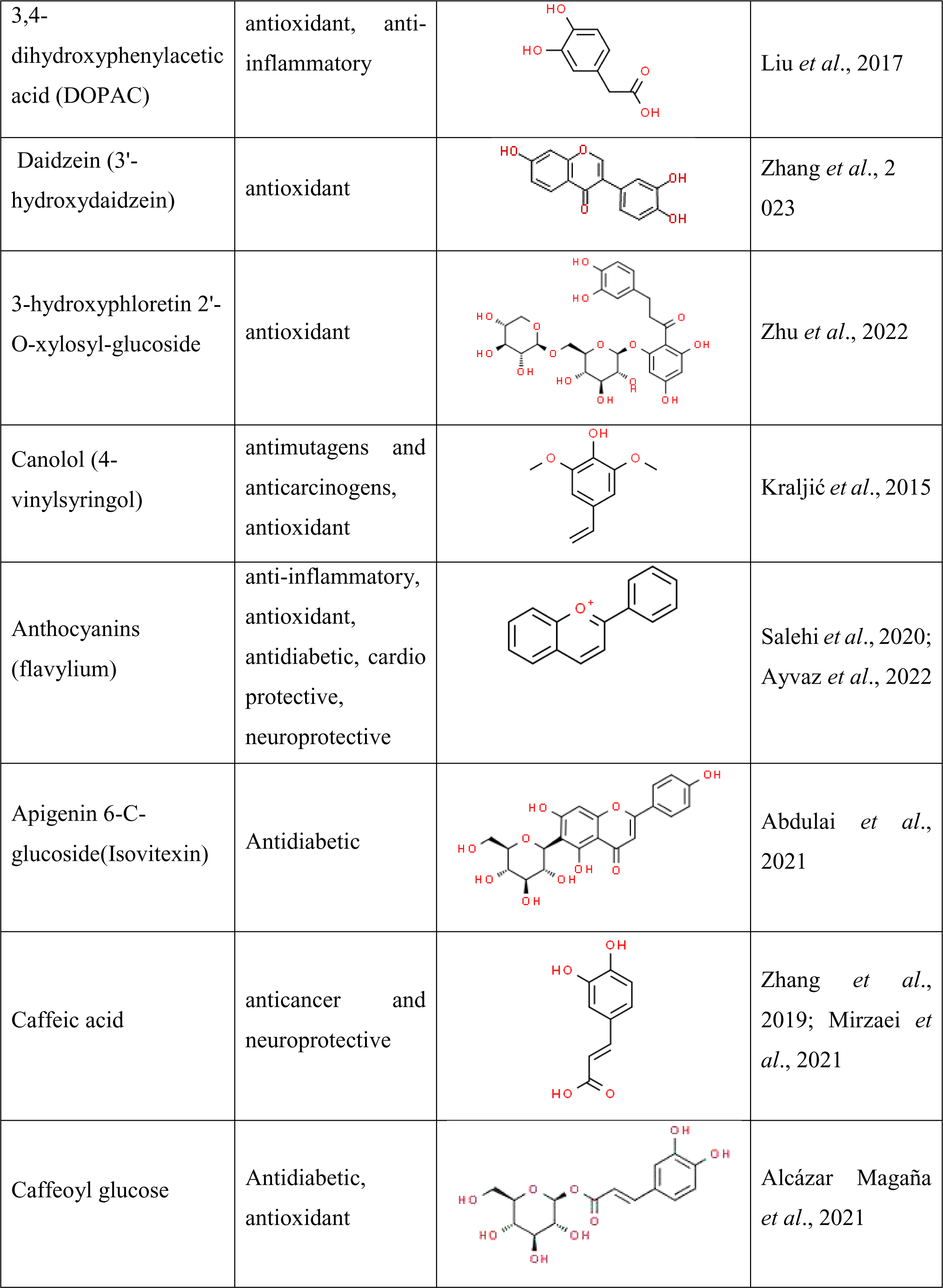
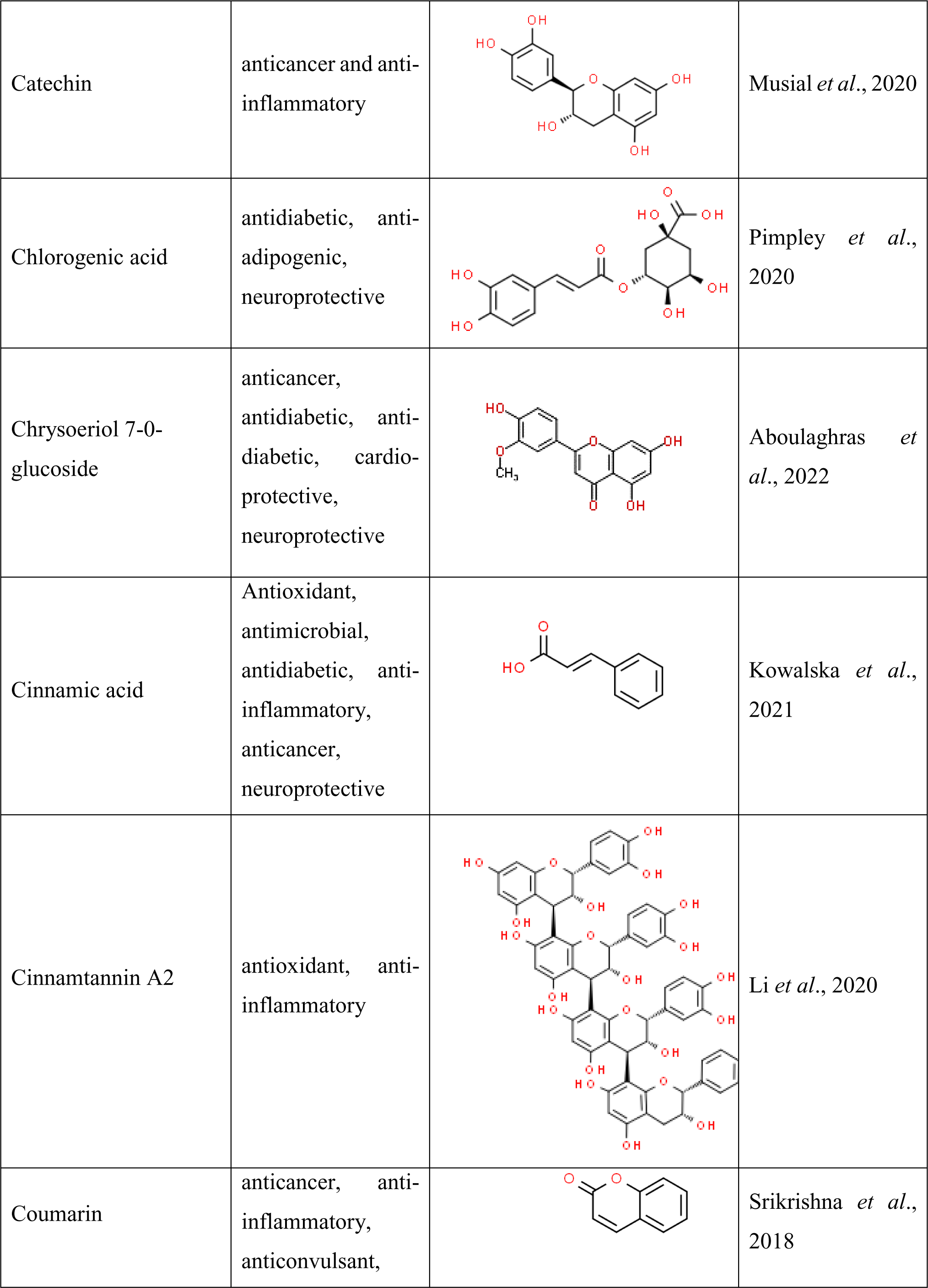
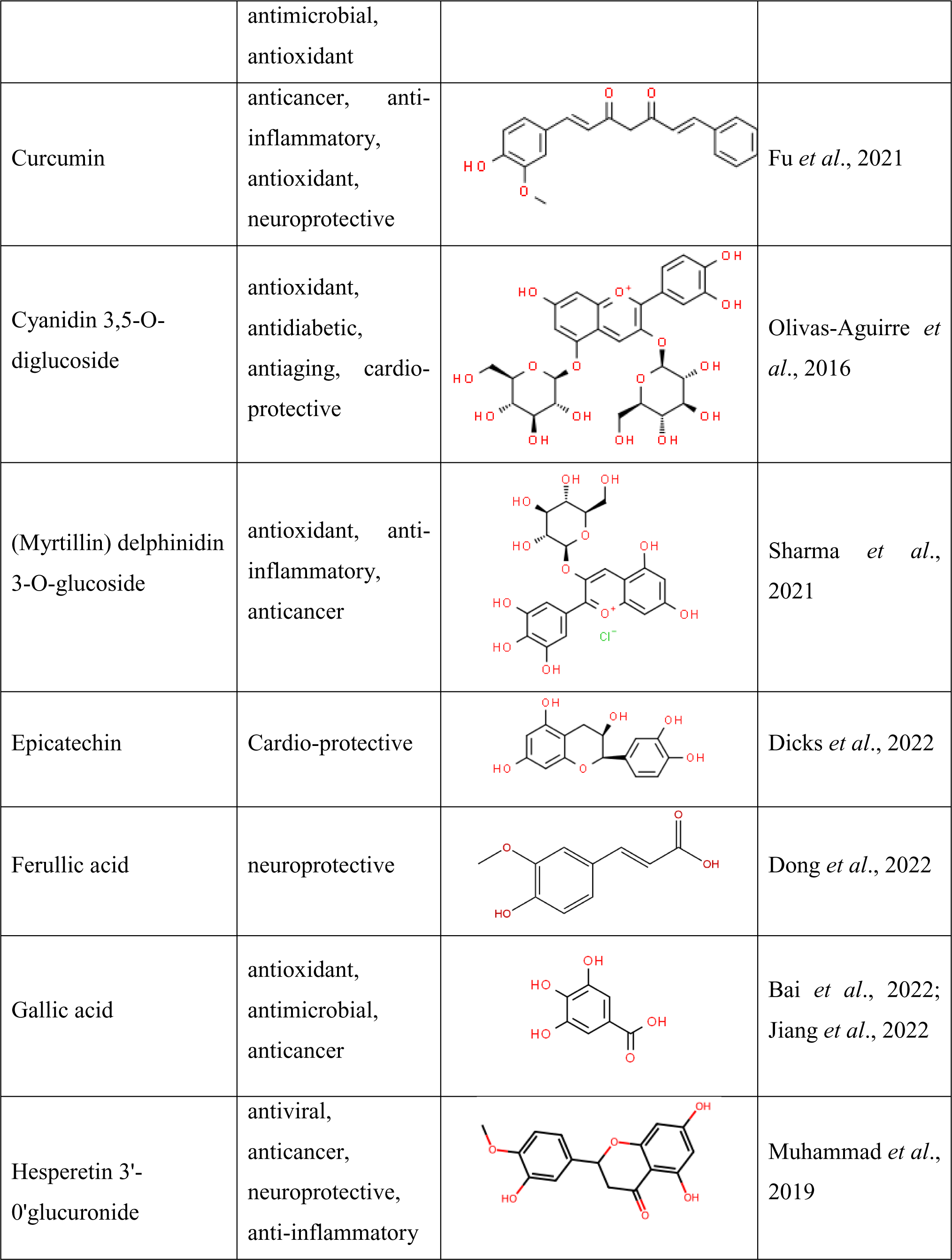
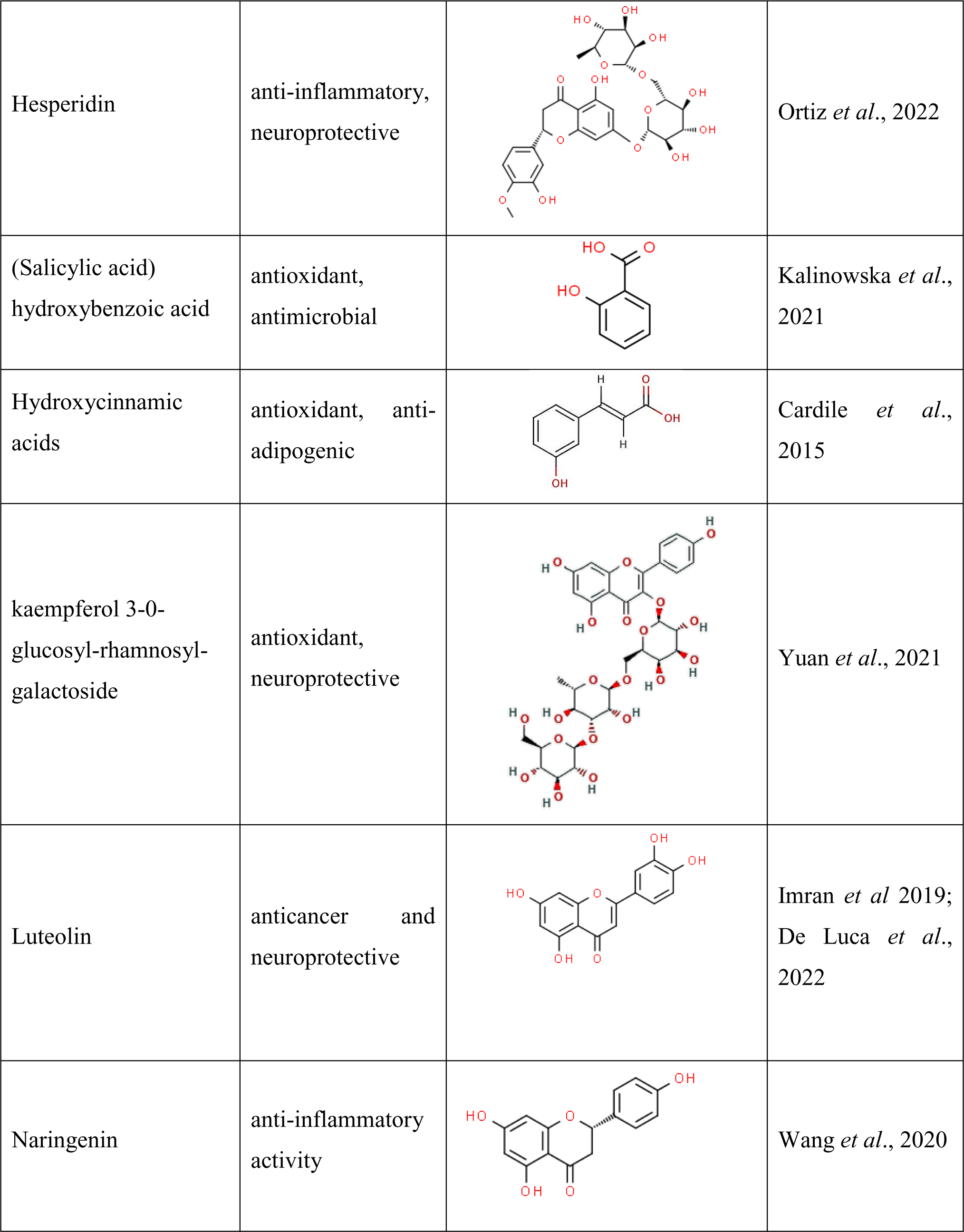
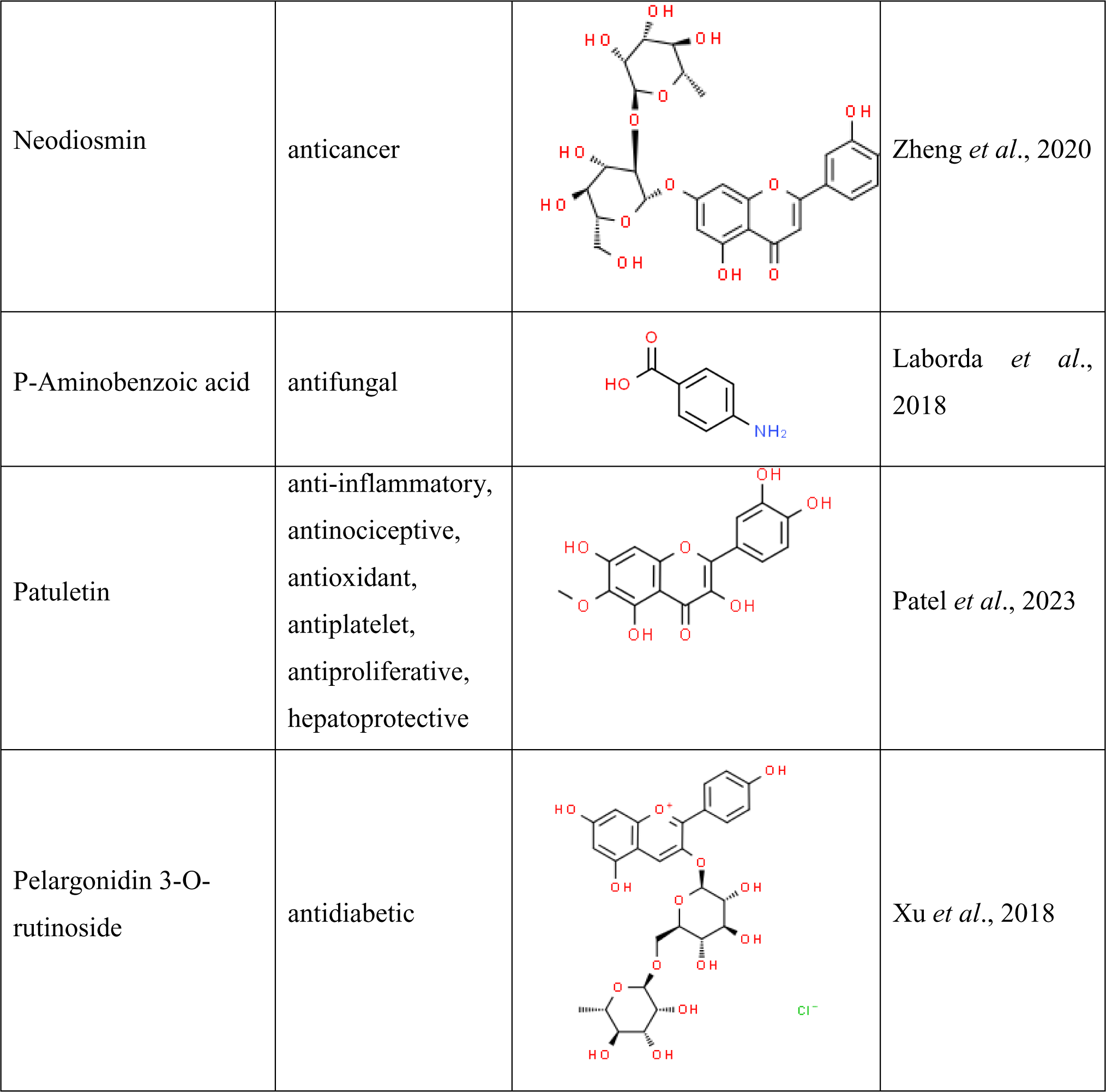
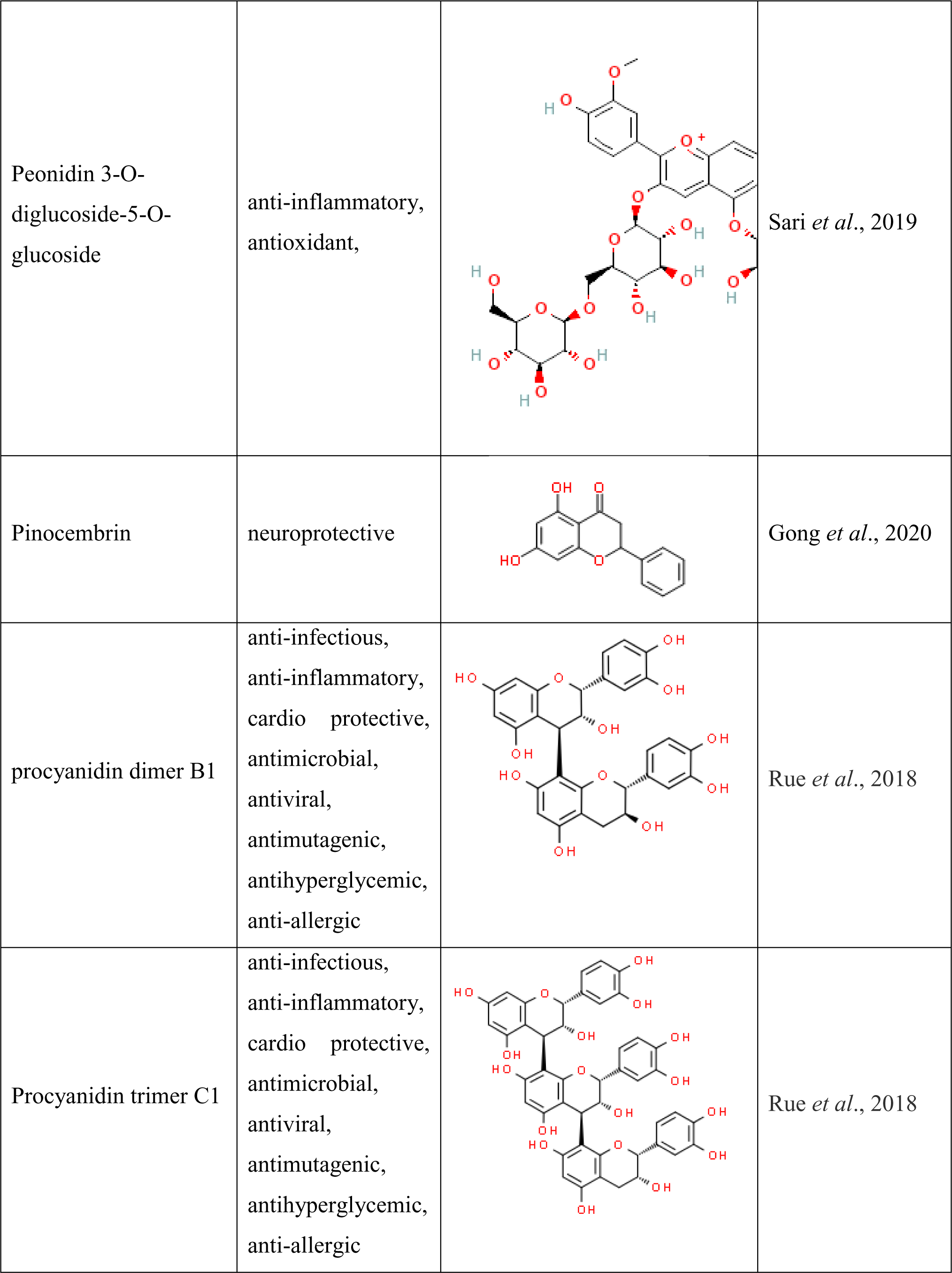
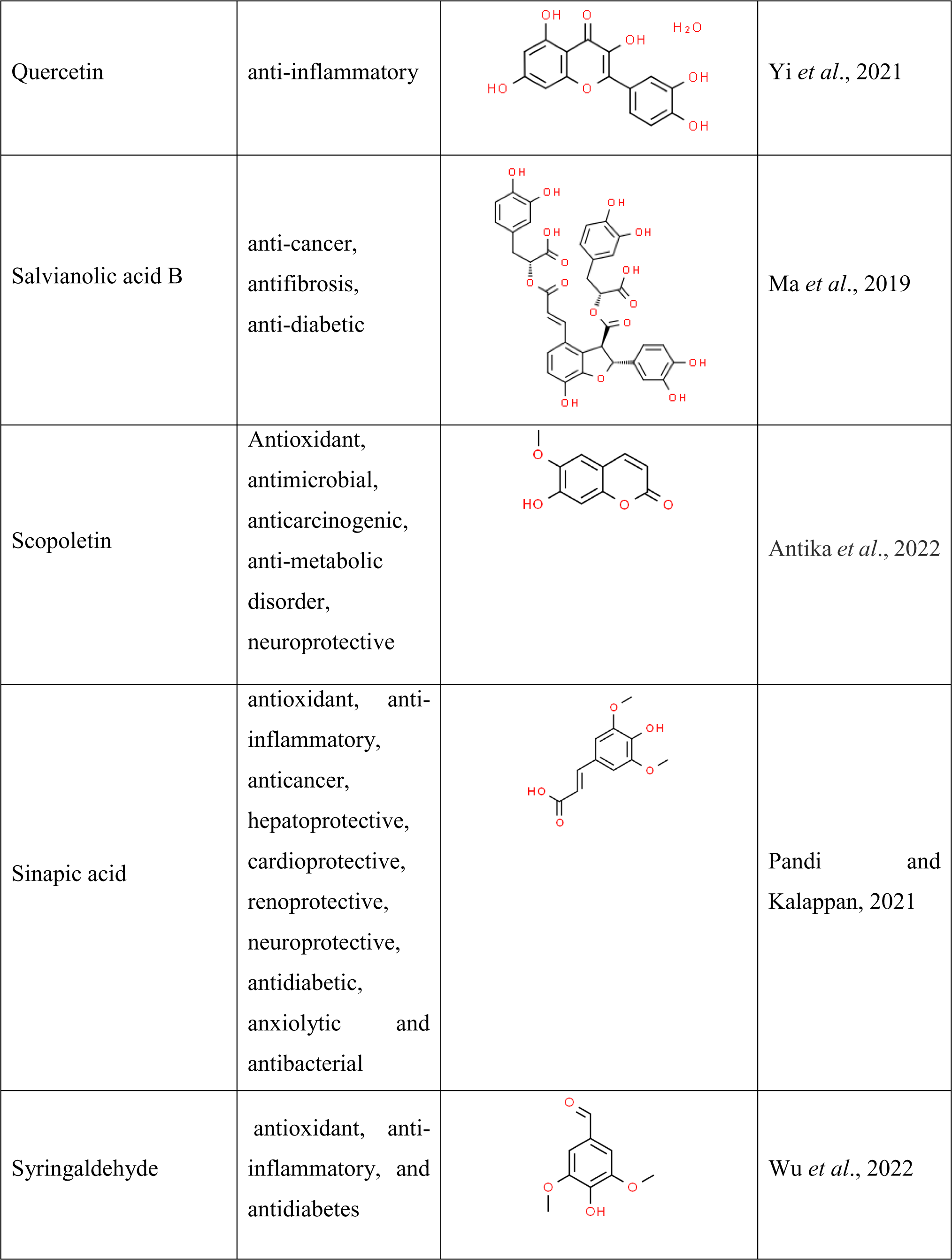
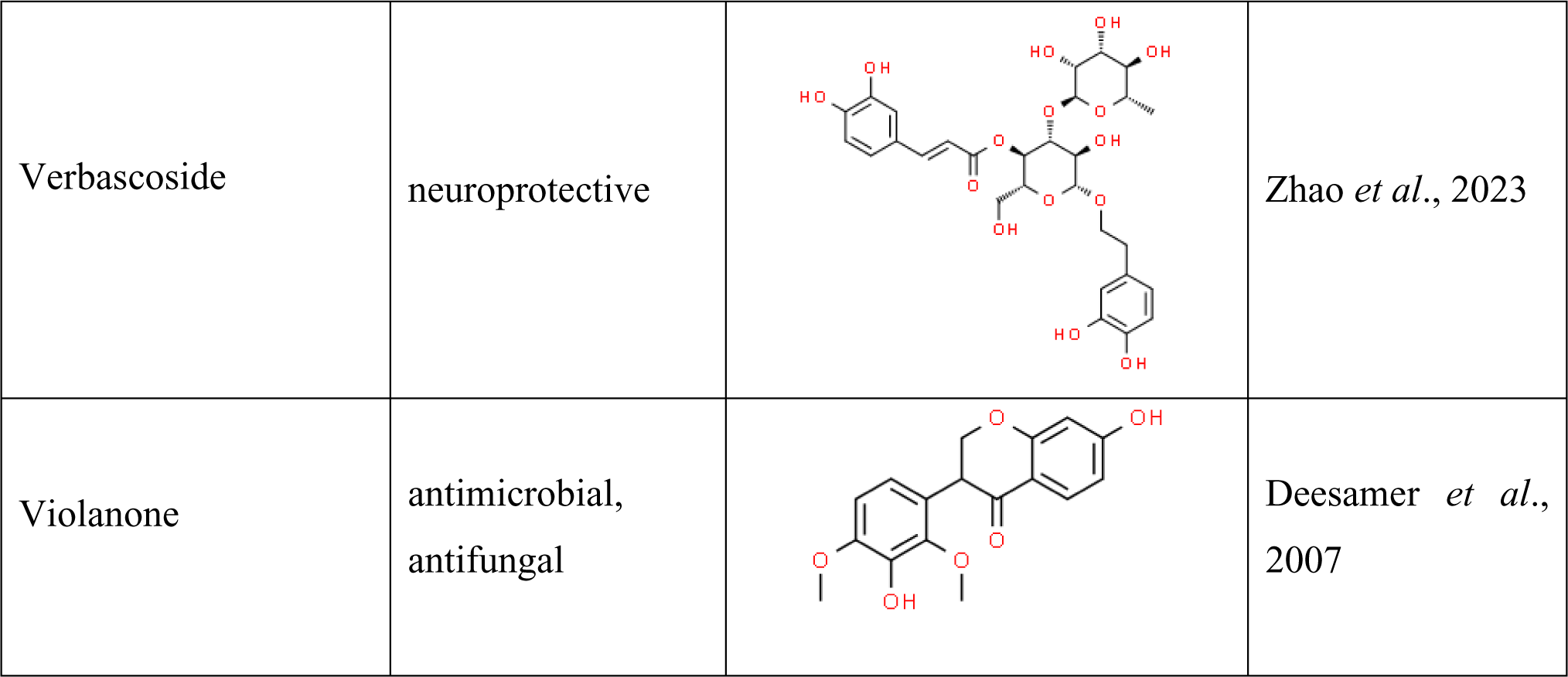
Pharmacological properties of major bioactive compounds identified in LC-MS, HPLC, and HPTLC analyses.

| Compound name | Pharmacological properties | Chemical Structure | References |
| --- | --- | --- | --- |
| 2-dehydro-O-desmethylangolensin | antioxidant |  | Ali *et al.*, 2021 |
| 3,4-dihydroxyphenylacetic acid (DOPAC) | antioxidant, anti-inflammatory |  | Liu *et al.*, 2017 |
| Daidzein (3'-hydroxydaidzein) | antioxidant |  | Zhang *et al.*, 2023 |
| 3-hydroxyphloretin 2'-O-xylosyl-glucoside | antioxidant |  | Zhu *et al.*, 2022 |
| Canolol (4-vinylsyringol) | antimutagens and anticarcinogens, antioxidant |  | Kraljić *et al.*, 2015 |
| Anthocyanins (flavylium) | anti-inflammatory, antioxidant, antidiabetic, cardio protective, neuroprotective |  | Salehi *et al.*, 2020; Ayvaz *et al.*, 2022 |
| Apigenin 6-C-glucoside(Isovitexin) | Antidiabetic |  | Abdulai *et al.*, 2021 |
| Caffeic acid | anticancer and neuroprotective |  | Zhang *et al.*, 2019; Mirzaei *et al.*, 2021 |
| Caffeoyl glucose | Antidiabetic, antioxidant |  | Alcázar Magaña *et al.*, 2021 |
| Catechin | anticancer and anti-inflammatory |  | Musial *et al.*, 2020 |
| Chlorogenic acid | antidiabetic, anti-adipogenic, neuroprotective |  | Pimpley *et al.*, 2020 |
| Chrysoeriol 7-O-glucoside | anticancer, antidiabetic, anti-diabetic, cardio-protective, neuroprotective |  | Aboulaghras *et al.*, 2022 |
| Cinnamic acid | Antioxidant, antimicrobial, antidiabetic, anti-inflammatory, anticancer, neuroprotective |  | Kowalska *et al.*, 2021 |
| Cinnamtannin A2 | antioxidant, anti-inflammatory |  | Li *et al.*, 2020 |
| Coumarin | anticancer, anti-inflammatory, anticonvulsant, antimicrobial,<br>antioxidant |  | Srikrishna *et al.*, 2018 |
| Curcumin | anticancer, anti-inflammatory,<br>antioxidant,<br>neuroprotective |  | Fu *et al.*, 2021 |
| Cyanidin 3,5-O-diglucoside | antioxidant,<br>antidiabetic,<br>antiaging, cardio-protective |  | Olivas-Aguirre *et al.*, 2016 |
| (Myrtillin) delphinidin 3-O-glucoside | antioxidant, anti-inflammatory,<br>anticancer |  | Sharma *et al.*, 2021 |
| Epicatechin | Cardio-protective |  | Dicks *et al.*, 2022 |
| Ferullic acid | neuroprotective |  | Dong *et al.*, 2022 |
| Gallic acid | antioxidant,<br>antimicrobial,<br>anticancer |  | Bai *et al.*, 2022;<br>Jiang *et al.*, 2022 |
| Hesperetin 3'-O-glucuronide | antiviral,<br>anticancer,<br>neuroprotective,<br>anti-inflammatory |  | Muhammad *et al.*, 2019 |
| Hesperidin | anti-inflammatory,<br>neuroprotective |  | Ortiz *et al.*, 2022 |
| (Salicylic acid)<br>hydroxybenzoic acid | antioxidant,<br>antimicrobial |  | Kalinowska *et al.*,<br>2021 |
| Hydroxycinnamic<br>acids | antioxidant, anti-adipogenic |  | Cardile *et al.*,<br>2015 |
| kaempferol 3-O-glucosyl-rhamnosyl-galactoside | antioxidant,<br>neuroprotective |  | Yuan *et al.*, 2021 |
| Luteolin | anticancer and<br>neuroprotective |  | Imran *et al* 2019;<br>De Luca *et al.*,<br>2022 |
| Naringenin | anti-inflammatory<br>activity |  | Wang *et al.*, 2020 |
| Neodiosmin | anticancer |  | Zheng *et al.*, 2020 |
| P-Aminobenzoic acid | antifungal |  | Laborda *et al.*, 2018 |
| Patuletin | anti-inflammatory,<br>antinociceptive,<br>antioxidant,<br>antiplatelet,<br>antiproliferative,<br>hepatoprotective |  | Patel *et al.*, 2023 |
| Pelargonidin 3-O-rutinoside | antidiabetic |  | Xu *et al.*, 2018 |
| Peonidin 3-O-diglucoside-5-O-glucoside | anti-inflammatory, antioxidant, |  | Sari *et al.*, 2019 |
| Pinocembrin | neuroprotective |  | Gong *et al.*, 2020 |
| procyanidin dimer B1 | anti-infectious, anti-inflammatory, cardio protective, antimicrobial, antiviral, antimutagenic, antihyperglycemic, anti-allergic |  | Rue *et al.*, 2018 |
| Procyanidin trimer C1 | anti-infectious, anti-inflammatory, cardio protective, antimicrobial, antiviral, antimutagenic, antihyperglycemic, anti-allergic |  | Rue *et al.*, 2018 |
| Quercetin | anti-inflammatory |  | Yi *et al.*, 2021 |
| Salvianolic acid B | anti-cancer,<br>antifibrosis,<br>anti-diabetic |  | Ma *et al.*, 2019 |
| Scopoletin | Antioxidant,<br>antimicrobial,<br>anticarcinogenic,<br>anti-metabolic<br>disorder,<br>neuroprotective |  | Antika *et al.*, 2022 |
| Sinapic acid | antioxidant, anti-inflammatory,<br>anticancer,<br>hepatoprotective,<br>cardioprotective,<br>renoprotective,<br>neuroprotective,<br>antidiabetic,<br>anxiolytic and<br>antibacterial |  | Pandi and<br>Kalappan, 2021 |
| Syringaldehyde | antioxidant, anti-inflammatory, and<br>antidiabetes |  | Wu *et al.*, 2022 |
| Verbascoside | neuroprotective |  | Zhao *et al.*, 2023 |
| Violanone | antimicrobial,<br>antifungal |  | Deesamer *et al.*,<br>2007 |

### 3.2. Identification and quantification of marker compounds by LC-MS

Detailed phytochemical profiling of hydroethanolic extract was performed using LC-MS analysis. LC-MS analysis showed a total of 22 phytochemicals, including 11 flavonoids, 2 polyphenols, 2 hydroxybenzoic acids, 4 hydroxycinnamic acid, 1 phenolic acid, and 2 phenolic aldehydes in a hydroethanolic fraction of *E. ganitrus* leaves, and the chromatogram was displayed in **Figure 4**. LC-MS data of the identified compounds with their retention time, responses (frequency), and concentration was provided in **Table 3**. The major eleven identified compounds were quercetin (803.0215 µg/L) at RT 7.319, Gallic acid (726.13 µg/L) at RT 6.223, Ferullic acid (652.34 µg/L) at RT 7.672, Chlorogenic acid (651.021 µg/L) at RT 8.812, Pinocembrin (264.11 µg/L) at RT 13.387, p-aminobenzoic acid (251.021 µg/L) at RT 1.678, Epicatechin (246.02 µg/L) at RT 1.336, Catechin (161.51 µg/L) at RT 1.336, Caffeic acid (123.31 µg/L) at RT 9.555, Syringaldehyde (116.31 µg/L) at RT 7.696, and Naringenin (106.31 µg/L) at RT 8.697 **(Figure 4 and 5)**.

**Figure 4.**
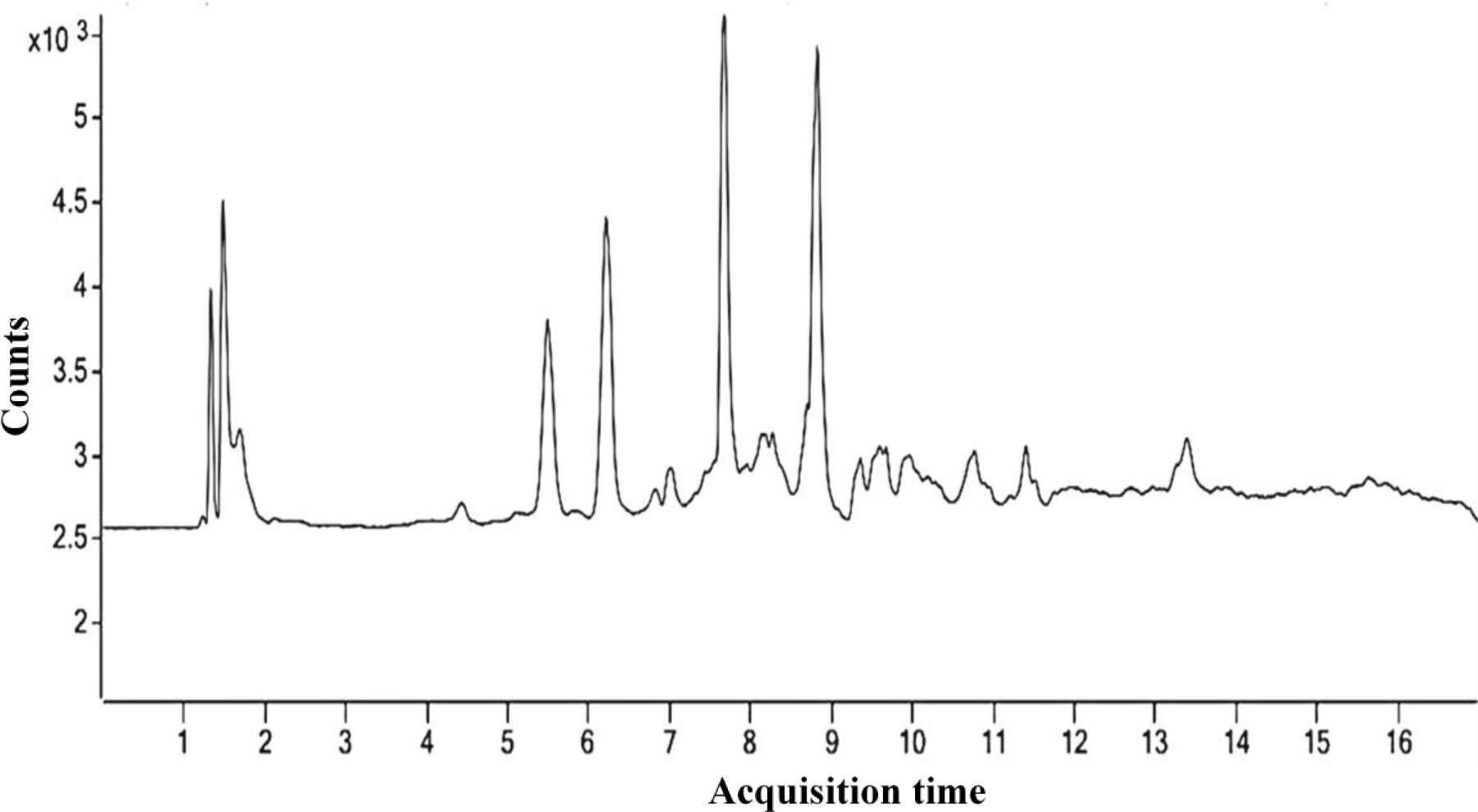
LC-MS chromatogram of identified phytochemical constituents’ profile hydrolethanolic extract of *E. ganitrus* leaves.

**Figure 5.**
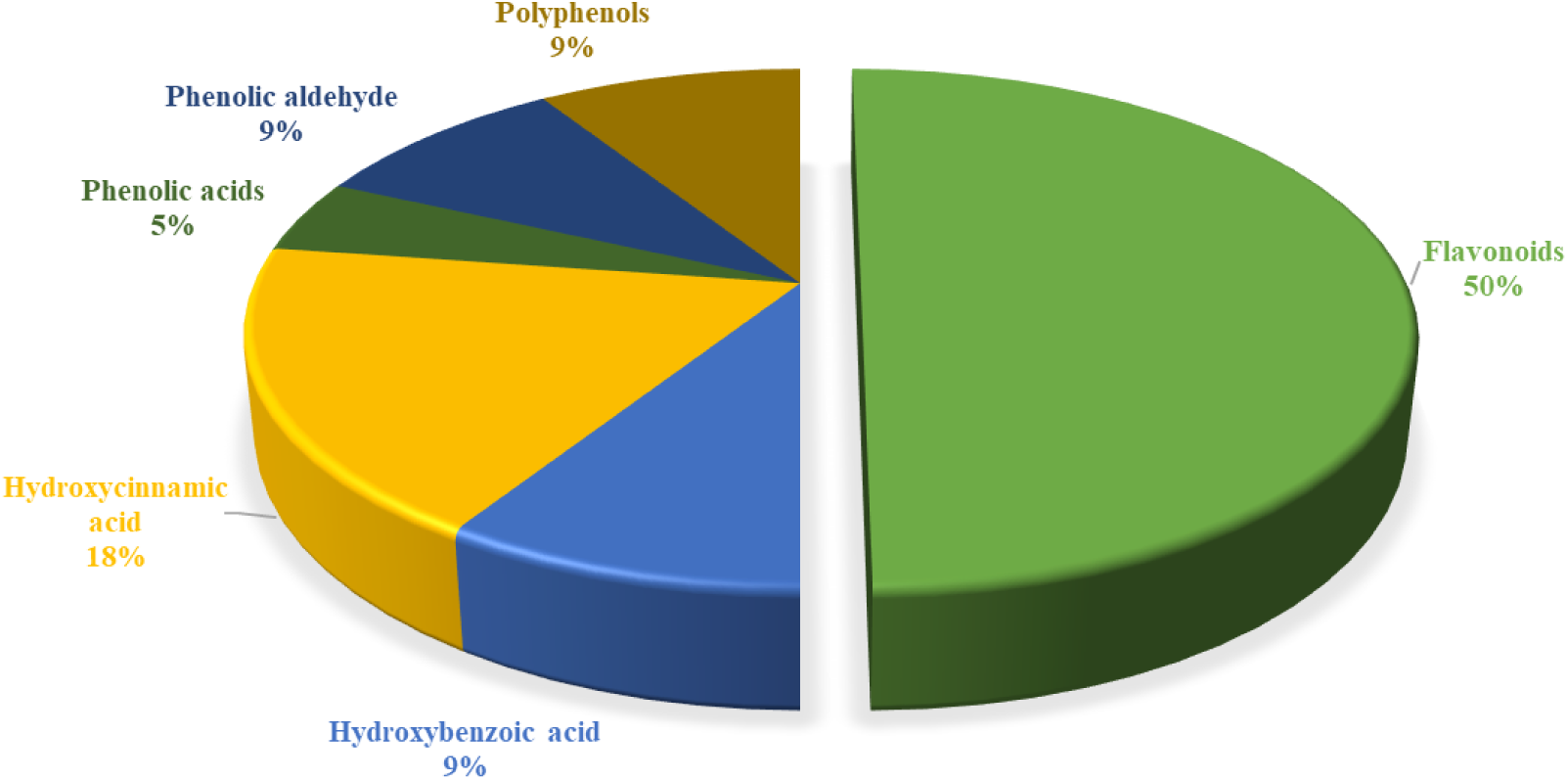
The phytochemical composition identified in LC-MS analysis in a hydroethanolic fraction of *E. ganitrus* leaves.

Identified compounds belonged to various classes, including flavonoids, polyphenols, hydroxybenzoic acid, hydroxycinnamic acid, phenolic acid, and phenolic aldehyde. The major bioactive compounds identified by LC-MS analysis were presented along with their classification and pharmacological activities **(Table 4)**. The bioactive compounds have diverse therapeutic potential which includes anti-inflammatory, antioxidant, antifungal, anticancer, antidiabetic, anti-adipogenic, cardio-protective and neuroprotective activities **(Laborda *et al*., 2018; Imran *et al*., 2019; Zhang *et al*., 2019; Musial *et al*., 2020; Gong *et al*., 2020; Pimpley *et al*., 2020; Yi *et al*., 2021; Mirzaei *et al*., 2021; Dong *et al*., 2022; Dicks *et al*., 2022; Bai *et al*., 2022; Jiang *et al*., 2022; Wu *et al*., 2022; De Luca *et al*., 2022)**.

#### HPTLC

The HPTLC analysis of the hydroethanolic extract of *E. ganitrus* leaves showed the presence of various phytoconstituents in different concentrations, such as Gallic acid (48.64 %), Curcumin (15.21 %), Caffeic acid (12.19%) and Cinnamic acid (6.50%) (**Figure 6**). The developed HPTLC method will assist in the standardization of *E. ganitrus* extract using biologically active chemical markers. Several pharmacological activities of identified phytochemicals exhibit antioxidant, anti-inflammatory, antifungal, and antibacterial activities due to the presence of bioactive phytochemicals such as phenolic acid, phenols, hydroxycinnamic acid, and flavonoids **(Zhang *et al*., 2019; Mirzaei *et al*., 2021; Fu *et al*., 2021; Bai *et al*., 2022; Jiang *et al*., 2022)**. The major bioactive compounds identified by HPTLC analysis were displayed with their classification and pharmacological activities in **Table 4**.

**Figure 6.**
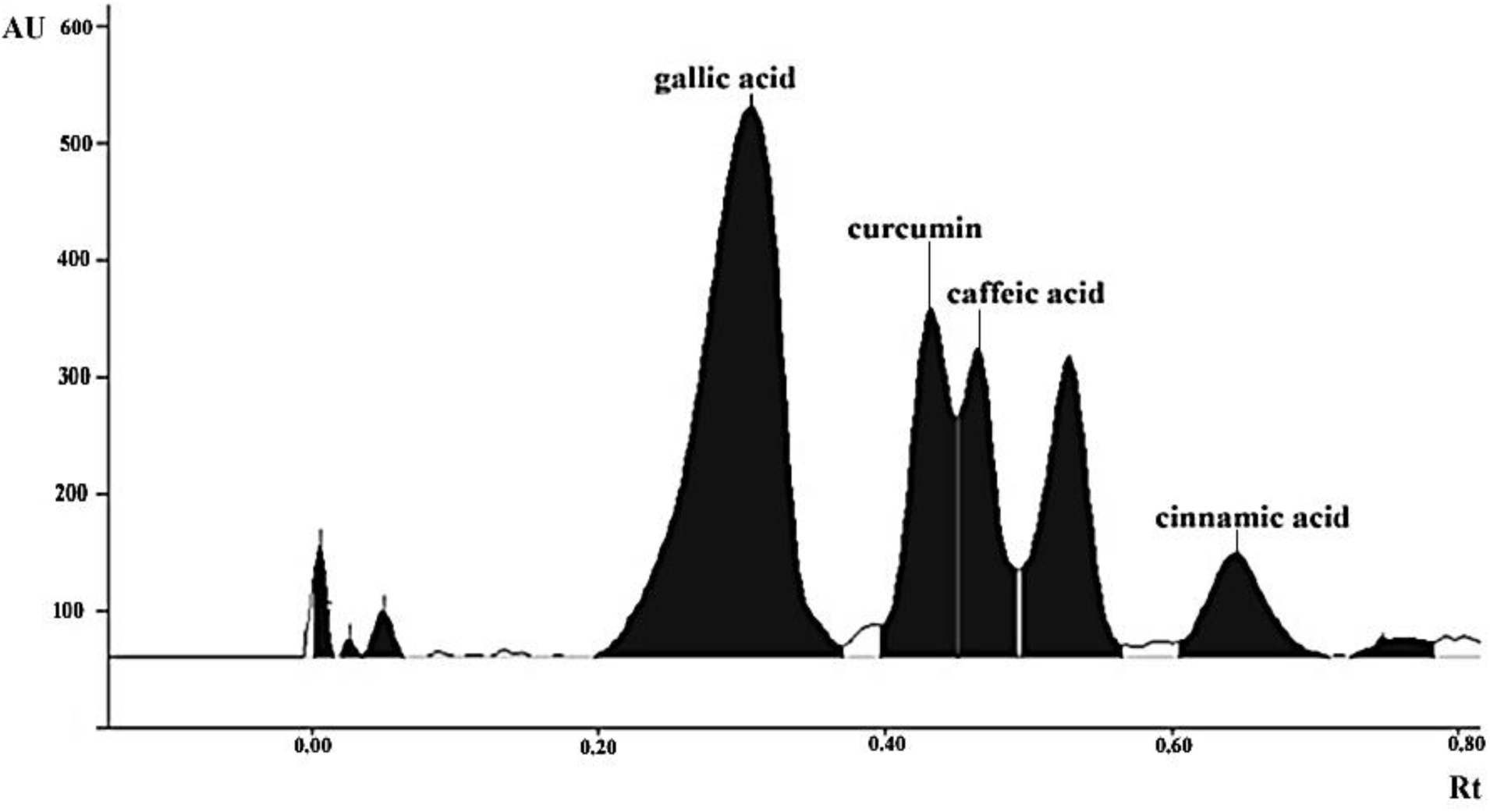
HPTLC chromatogram of identified phytochemical constituents’ profile hydroethanolic extract of *E. ganitrus* leaves.

#### Conclusion

The phytochemical profile of *E. ganitrus* leaf extract was characterized using HPLC, LC-MS, and HPTLC analyses. The hydroethanolic fraction of *E. ganitrus* leaf was found to contain valuable metabolites: phenolic acid, polyphenols, flavonoids, phenols, phenolic glycosides, flavonoid glycosides, terpene glycoside, phenylpropanoid glycoside, hydroxycinnamic acid, hydroxybenzoic acid, phenolic aldehyde, lignin, and tannins. The phytochemicals could be employed as potential biochemical markers because different phytochemicals were detected in three studied (HPLC, LC-MS, and HPTLC) analytical techniques. Previous research has shown that *Elaeocarpus* species contain beneficial bioactive compounds in significant amounts, which have a wide range of applications in the pharmaceutical, food, and cosmetic industries. Moreover, only a limited number of researches are available on the phytochemical profiling of *E. ganitrus.* Hence, extensive investigations on phytochemical analysis and pharmacological activities of different fractions of leaf, fruit (bead), and pulp of *E. ganitrus* would be of much interest using a combination of modern analytical techniques or assays. Further research is required to isolate and characterize individual bioactive compounds and to validate their therapeutic potential.

## Acknowledgment

The authors acknowledge all the faculty and staff members of the Department of Biotechnology, Department of Biomedical Engineering, Department of Agriculture Technology & Agri– Informatics, Shobhit Institute of Engineering & Technology, (Deemed-to-be University), Meerut, 250110, India for their support and motivation.

